# *In vivo* optical induction of I_NaL_ by photoactivation of AaH-II peptide in beating heart using wireless implantable device

**DOI:** 10.64898/2026.10.27.761894

**Authors:** Matthias Dereli, Hugo Millet, Agnès Hivonnait, Antoine Persello, Maureen Choteau-Bodor, Virginie Aillerie, Pascal Aumond, Claude Zoukimian, Edith Bigot, Remy Beroud, Romain Capoulade, Philipp Gutruf, Flavien Charpentier, Michel De Waard, Jérôme Montnach

## Abstract

Precise spatiotemporal control of pharmacological compounds in cardiac tissue remains an unmet challenge. Systemic drug administration poorly reproduces the localized and transient nature of cardiac events. Photopharmacology, in which light controls drug activity, offers a compelling solution, but its translation to the beating heart requires wireless light delivery to a moving, soft, and electrically active organ. Here, we demonstrate the first in vivo use of a wireless device to photoactivate a peptide that modulates cardiac excitability. Using a flexible, battery-free implantable optoelectronic device, we achieved light-triggered uncaging of AaH-II*, a natural peptide that enhances the late sodium current. We first validated the caged peptide in vitro and ex vivo, showing that caging suppresses its activity and that UV illumination rapidly restores it. In vivo, wireless photoactivation produced a clear modulation of cardiac electrical activity, as monitored by electrocardiography. This work establishes device-based peptide photomodulation as a strategy for remote, temporally controlled pharmacological intervention in the heart. More broadly, it provides a new platform for probing cardiac excitability and developing region-targeted therapies.

## Introduction

Precise spatiotemporal control of pharmacological compounds in living tissues remains a fundamental challenge in cardiovascular research. Current drug delivery strategies rely almost exclusively on systemic administration, resulting in widespread cardiac and extracardiac effects that address the localized and transient nature of many cardiac events, such as arrhythmias ^1^. Most of the ion channels targeted by antiarrhythmic drugs are broadly expressed across cardiac cell types and extra-cardiac tissues ^2^. Also, the pharmacokinetic profiles of drugs are often characterized by slow absorption & tissue distribution equilibrium, complex metabolism and progressive excretion spanning hours to days ^3^ that prevent on-demand activation and rapid termination of drug action. Together, these spatial and temporal limitations impose a hard ceiling on the precision achievable with conventional pharmacology.

Reflecting these constraints, major therapeutic advances over the past two decades have shifted toward interventional strategies. Implantable pacemakers, cardioverter-defibrillators (ICDs) ^4^, and catheter ablation ^5^ offer the on-demand, cardiac-restricted precision that pharmacology lacks. However, they remain invasive and technically demanding, with risks of mechanical mismatch, collateral damage, and procedure failure ^6–8^. Also, none of the technologies rely on pharmacology and may lack the mechanistic refinements that drugs offer.

Photopharmacology, defined as a set of technologies for light-mediated regulation of drug activity, has emerged as a strategy to bridge this gap. Drug activity can be switched on, off, or fine-tuned with light, granting unprecedented spatiotemporal control over compound delivery ^9,10^. Unlike cardiac optogenetics ^11–15^, most photopharmacology approaches do not require genetic modification, and rely instead on photoactivatable or photoswitchable molecules that can be locally uncaged or isomerized, respectively, at the desired time and site. Despite striking successes for vision restoration ^16,17^, nociception ^18^ or pain modulation ^19^, cardiac applications have so far remained limited ^20,21^. We recently developed a photopharmacology approach that exploits the ion channel targeting properties of natural peptides to modulate cell excitability *in vivo* upon application of light ^22,23^. Strategic grafting of a UV-photocleavable o-nitroveratryloxycarbonyl (Nvoc) protecting group onto a critical amino acid residue renders the peptide inactive until light triggers the group cleavage and the active peptide uncaging. This photoactivation technology enables highly localized, on-demand delivery of active peptide and was validated across *in vitro*, *ex vivo*, and *in vivo* models of neuronal and cardiac activity ^23^. However, translating this strategy *in vivo* to the beating heart requires light-delivery hardware compatible with a moving, soft and electrically active organ - a constraint that conventional optical fibers and tethered cannulas cannot meet ^24–26^. Recent advances in miniaturized, battery-free, flexible, and fully implantable optoelectronic devices now enable untethered and programmable tissue illumination ^7,8,19,27,28^, providing the missing hardware layer for investigating if cardiac peptide photopharmacology is a viable strategy. For this first proof-of-concept, we used a caged version of AaH-II*, a peptide that was previously carefully investigated for its remarkable proarrhythmic cardiovascular effects through the induction of I_NaL_ ^29^. This strategy eases significantly the demonstration that controlled photoactivation of a caged peptide can reliably alter cardiac electrical activity by producing a clear, monitorable electrophysiological signature ^29^.

Here, we report the first wireless and device-mediated *in vivo* cardiac photoactivation of a caged peptide that leads to cardiac excitability modification. We describe the synthesis and functional validation of caged AaH-II* *in vitro* and *ex vivo*, and provide the first *in vivo* evidence that device-triggered local photoactivation can fine-tune cardiac electrical activity. This work establishes a platform for spatiotemporally controlled compound delivery to the heart, with broad implications for probing cardiac excitability and, ultimately, for developing region-targeted pharmacological strategies.

## Results

Molecular engineering of photoactivatable sodium channel activator - To demonstrate the potential of device-mediated *in vivo* peptide-based photopharmacology, we used AaH-II, a 64-aminoacid peptide that slows sodium channel inactivation and leads to an easily distinguishable arrhythmic phenotype. To produce an inactive AaH-II peptide that is sensitive to light, we chose to chemically synthetize AaH-II* with a photosensitive orthogonal o-nitroveratryloxycarbonyl (Nvoc) cage group (**Fig. 1a**), as previously reported ^22,23^. For the position of the Nvoc group, R^62^ was chosen because of its importance for the interaction of AaH-II with the chimeric Na_V_1.7-Na_V_PaS channel ^30^. R^62^ acts as a molecular anchor by interacting simultaneously with Q^265^ from the pore region of domain I and E^1589^ of the S3-S4 loop of domain IV (**Fig. 1a**). To graft the Nvoc group we first replaced arginine 62 residue by lysine (AaH-II-R^62^K, termed herein AaH-II*). We previously reported that this substitution does not alter the pharmacological properties of the peptide ^29^. We established that the addition of the Nvoc group on the lateral chain of K^62^ induces severe van der Walls steric clashes with A^161^ and Y^224^ of Na_V_1.7-Na_V_PaS. Nvoc addition onto K^62^ should be sufficient to drastically reduce the peptide potency onto voltage-gated sodium channels (**Fig. 1b**). Caged AaH-II* was assembled stepwise using Fmoc-based solid phase peptide synthesis (**Supplementary Fig. 1a-c**). Proper folding of caged peptide was confirmed by a shift in the elution profile to greater water polarity. The proper mass of caged AaH-II* was confirmed with a measured *m/z* ratio of 828.7923 [M+9H]^9+^ corresponding to a peptide of 7450.13 Da (theoretical monoisotopic mass of 7449.18 Da), (**Supplementary Fig. 1d**) indicating proper peptide assembly and folding, and the preservation of the Nvoc group during the entire synthesis process (non-caged AaH-II* monoisotopic mass of 7210.14 Da for comparison). Compared to non-caged AaH-II*, caged AaH-II* presents an elution profile that is less polar on RP-HPLC, consistent with hydrophobic Nvoc properties (**Fig. 1c**). Nvoc photolysis is reportedly most efficient at 350 and 365 nm ^23,31^. Illumination of caged AaH-II* at 365 nm produces uncaged AaH-II* with an uncaging half-time of 1.6 min (45 mW/cm^2^) and half-power of 9.2 mW/cm^2^ (**Supplementary Fig. 2**). Photolysis of caged AaH-II* efficiently also occurred at 405 nm (88.7% of uncaging) but became negligible at 435-740 nm wavelengths (10 min, irradiation power > 18 mW/cm^2^; **Supplementary Fig. 3**). Co-elution experiments of non-caged AaH-II* with fully uncaged AaH-II* (after 30 min illumination of caged AaH-II* at 365 nm and 45 mW/cm²) results in a single elution peak suggesting that Nvoc photolysis releases a peptide fully similar to non-caged AaH-II* (**Fig. 1d**). This result was confirmed by mass spectrometry analyses indicating similar molecular mass (uncaged AaH-II*: *m/z* = 902.2720 [M+8H]^8+^ corresponding to a peptide of 7210.16 Da, similar to non-caged AaH-II*: *m/z* = 802.1345 [M+9H]^9+^ corresponding to a peptide of 7210.21 Da).

**Figure 1.**
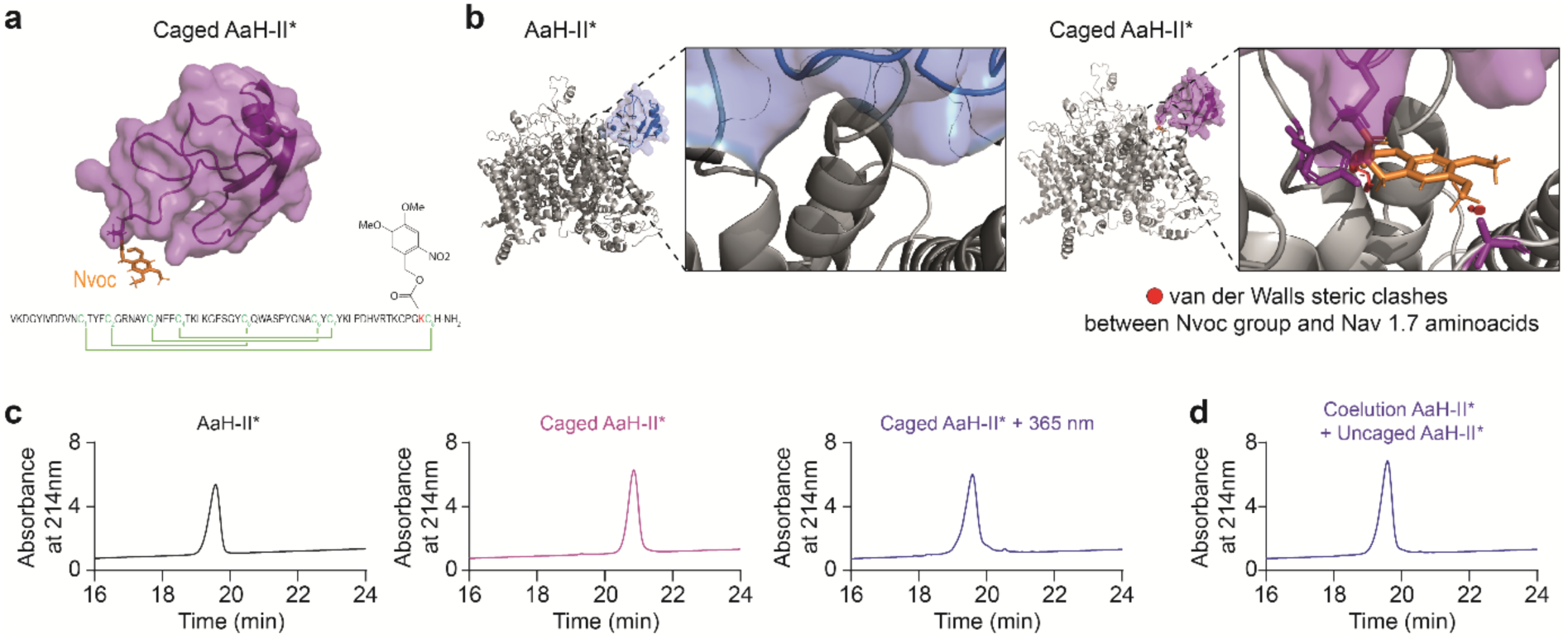
Engineering of photoactivatable AaH-II* for optical modulation of sodium channels. (a) Structural modeling and amino acid sequence of AaH-II*-Nvoc, with the R^62^K substitution and Nvoc group highlighted in orange, respectively. (b) Modeling of native AaH-II* (left) and caged AaH-II* (right) docked onto the chimeric Na_V_1.7-Na_V_PaS domain. Van der Waals steric clashes between the Nvoc group and Na_V_1.7-Na_V_PaS amino acid residues are highlighted in red. (c) RP-HPLC elution profiles of native AaH-II* (left panel), caged AaH-II* (middle panel), and uncaged AaH-II* (right panel). (d) Coelution profile of combined native AaH-II* and fully uncaged AaH-II*. Source data are provided as a Source Data File.

Light-induced activation of sodium channels - Using a high-throughput patch-clamp system equipped with an illumination system, we confirmed that caged AaH-II* had no impact on hNa_v_1.2 inactivation kinetic within a range of concentrations (0.01 to 1 nM) at which non-caged AaH-II* displays potent effects as previously reported (**Fig. 2a-c**) ^29^. For instance, 1 nM caged AaH-II* produced a minimal 1.7-fold increase of the area under the curve (AUC), relative to untreated hNa_v_1.2 channels. In contrast, application of 1 nM of uncaged AaH-II* produced a significant 9.5-fold increase of the area under the curve (AUC). Concentration-response curves illustrate that photoactivation of AaH-II* significantly increased its potency by 63-fold (caged AaH-II*: EC_50_ = 99.8 nM (n = 77 cells) *versus* uncaged AaH-II*: EC_50_ = 1.58 nM (n = 99 cells)) (**Fig. 2b**). As a result of this finding, application of 1 nM of caged AaH-II* had no effect on sodium current properties, whereas a 3-min illumination at 365 nm (45 mW/cm^2^) induced a progressive and significant slowing of hNa_v_1.2 sodium current inactivation and hence an increase in the AUC (AUC = 4.4 ± 0.4 at 365 nm *versus* 1.02 ± 0.03 in dark, n=15 cells, p<0.0001) (**Fig. 2d-f**).

**Figure 2.**
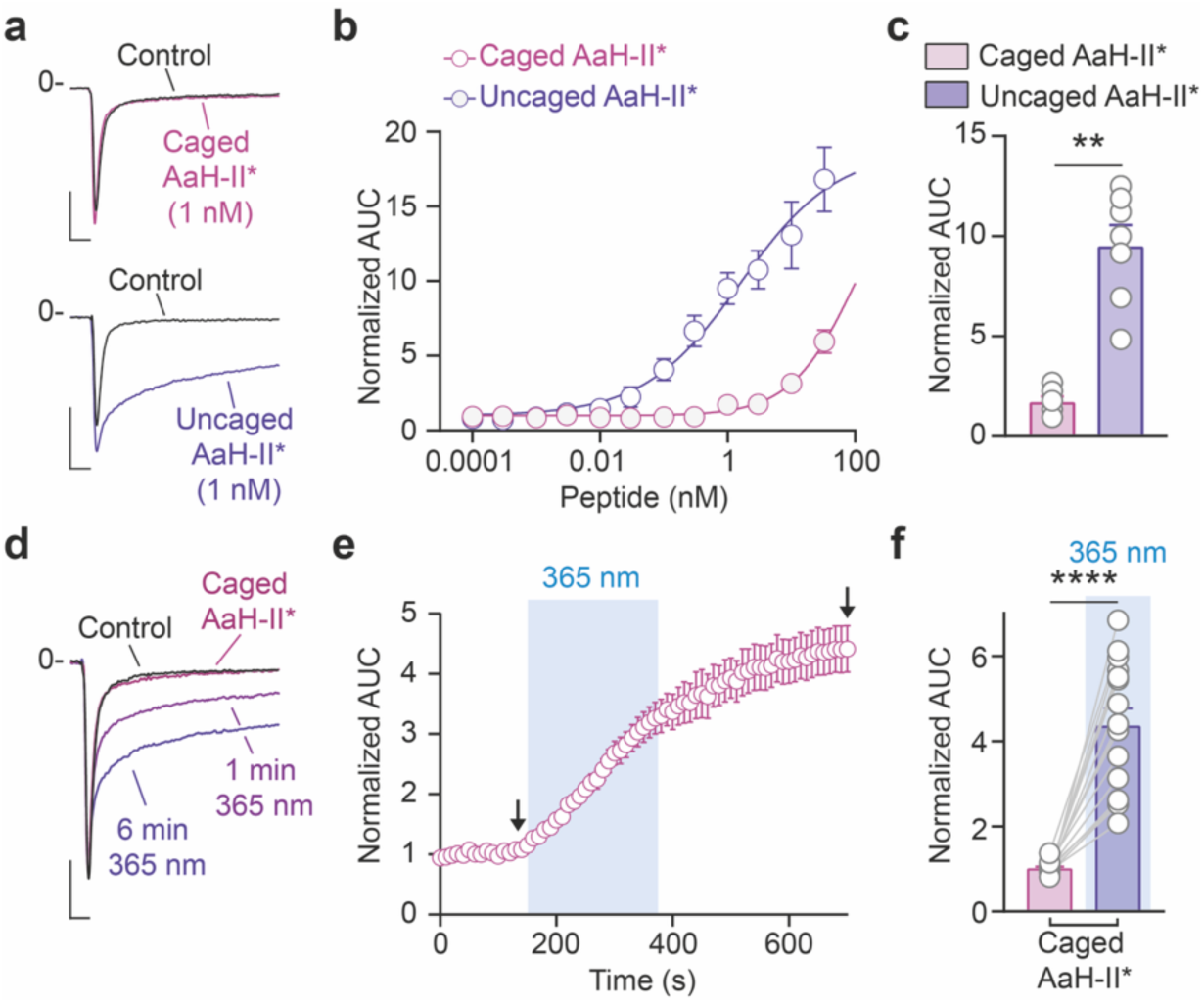
Optical activation of caged AaH-II* induces persistent sodium current. (a) Representative hNa_V_1.2-mediated sodium current traces recorded in the presence of 1 nM caged AaH-II* (top panel) and uncaged AaH-II* (bottom panel). Scale bars = 2 ms, 1000 pA. (b) Concentration-response curves of caged AaH-II* (purple) and uncaged AaH-II* (navy blue) on hNa_v_1.2 channels (n = 3-12). (c) AUC quantification of the effects of 1 nM caged AaH-II* (n = 6) and uncaged AaH-II* (n = 7) on hNa_v_1.2-mediated current. (d) Representative sodium current traces recorded under control conditions (black), in the presence of 1 nM caged AaH-II* (purple), and after 1 and 6 min of 365 nm illumination (blue). Scale bars = 2 ms, 1000 pA. (e) Time course of the light-induced increase in AUC during photoactivation of caged AaH-II* (n = 15). (f) Quantification of photoactivation kinetics and the resulting effects of 1 nM caged AaH-II* (n = 6) and uncaged AaH-II* (n = 7) on hNa_v_1.2-mediated current. Data are presented as mean ± SEM. Statistical analysis: panel c, Friedman test followed by Dunn’s multiple-comparison test *versus* baseline; panel f, Wilcoxon matched-pairs signed-rank test. Statistical significance is indicated as **, p < 0.01 and ****, p < 0.0001. Source data are provided as a Source Data File.

Light-induced activation of sodium channels - The effects of caged AaH-II* photoactivation were evaluated in isolated rat heart perfused in Langendorff mode by illuminating the epicardial surface of the right ventricle (**Fig. 3a**). Perfusion of 10 nM caged AaH-II* or its photoactivation has no impact on interbeat interval (**Fig. 3b-c**) or on cardiac conduction within the illuminated region **(Supplementary Fig. 4a-b),** suggesting that effects were restricted to the illuminated area and that Na_v_1.5 is not affected. Based on the reported effects of increased late sodium current due to slowing of inactivation of sodium channels, we expected to observe ventricular repolarization perturbations. Perfusion of 10 nM caged AaH-II* has no effect on either the duration nor the amplitude of the repolarization wave of the recorded area (repolarization wave duration: 53.57 ± 8.38 ms during 10 nM caged AaH-II* perfusion *versus* 46.49 ± 6.41 ms at baseline; repolarization wave area: 5.8 ± 21.5% increase relative to baseline; n=5 hearts), while 365 nm illumination led to a progressive increase of the repolarization wave duration (from 46.5 ± 6.4 ms at baseline to 110.5 ± 15.7 ms ms after 9 min of 365 nm, p<0.05) and of the repolarization wave area (108.5 ± 44.7 % increase relative to the baseline after 9 min of 365 nm illumination, p<0.05 for both) (**Fig. 3b-f**). In addition, photoactivation of 10 nM caged AaH-II* induces evident signs of arrhythmias as witnessed by the number of hearts (4/6) presenting premature ventricular beats and an alteration of the homogeneity of the cardiac propagation within the recorded area (**Fig. 3g-i**).

**Figure 3.**
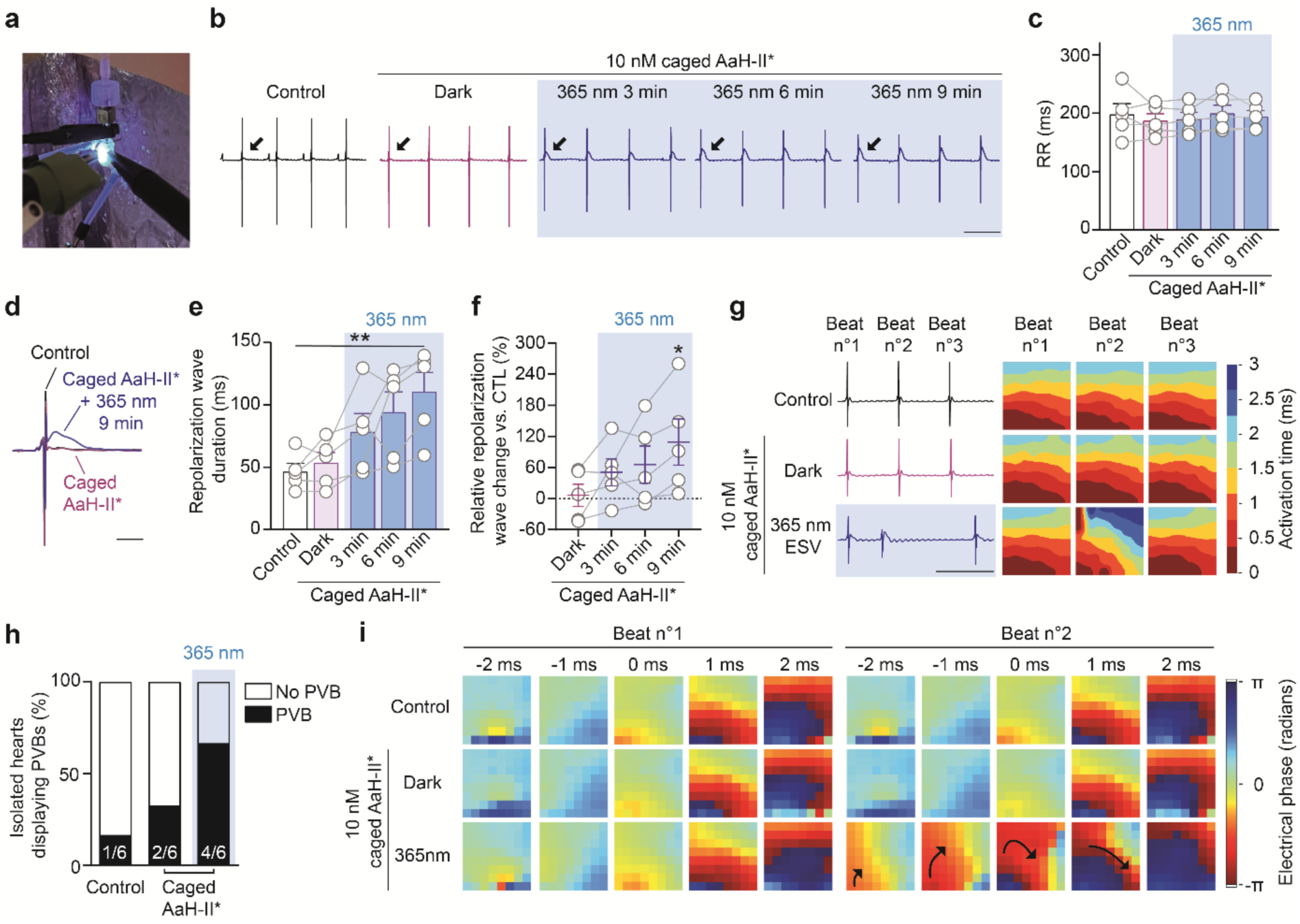
Photoactivation of caged AaH-II alters ventricular repolarization. (a) Image of an isolated Langendorff-perfused rat heart positioned under the MEA recording system, with the illuminated region indicated in blue. (b) Representative electrical field potential traces recorded under control conditions (black), during perfusion with 10 nM caged AaH-II* in the dark (purple), and at 3, 6 and 9 min of 365 nm illumination (blue). Scale bar = 200 ms. (c) Quantification of RR intervals measured under the same experimental conditions as the representative traces (n = 5). (d) Representative electrical field potential traces illustrating the increase in repolarization wave area and duration under control conditions, during perfusion with 10 nM caged AaH-II* in the dark, and after 9 min of 365 nm illumination. Scale bar = 20 ms. (e) Quantification of repolarization wave duration measured under the same experimental conditions as the representative traces (n = 5). (f) Relative fold change in repolarization wave area normalized to control, measured under the same experimental conditions as the representative traces (n = 5). (g) Representative MEA traces under control conditions, during 10 nM caged AaH-II* perfusion in the dark and 9 min of 365 nm illumination showing ectopic ventricular stimulation at beat 2 (left panel), with corresponding activation maps (right panel). Scale bar = 200 ms. (h) Incidence of premature ventricular beats during perfusion with 10 nM caged AaH-II* under right ventricular epicardial illumination (n = 6 hearts). (i) Representative phase maps during control, perfusion with 10 nM caged AaH-II* in the dark and during 365 nm illumination highlighting the arrhythmogenic impact of activated AaH-II* on the directionality of the electrical propagation (black arrow). Data are presented as mean ± SEM. Statistical analysis: panels c and e, Friedman test followed by Dunn’s multiple-comparison test *versus* baseline; panel f, Friedman test followed by Dunn’s multiple-comparison test *versus* baseline using repolarization wave area data. Statistical significance is indicated as *, p < 0.05 and **, p < 0.01. Source data are provided as a Source Data File.

To demonstrate the feasibility of *in vivo* cardiac photopharmacology, we used an anesthetized rat model of right-mini-thoracotomy (**Fig. 4a**). Intravenous injection of 33 µg/kg of caged AaH-II* had no impact on cardiac electrical activity. However, fiber-mediated illumination of the right ventricle significantly increases the area of the T wave (+23.8 ± 10.5 % relative to control period, p<0.05, n= 7 rats) without affecting QTc interval (66.7 ± 1.2 ms after illumination *versus* 65.2 ± 1.1 ms during baseline period, n= 7 rats). Importantly, we also reported that other ECG parameters, such as RR and QRS intervals, were not affected by AaH-II* photoactivation (RR interval: 145.7 ± 3.6 ms after illumination *versus* 148.2 ± 3.9 ms at baseline; QRS interval: 18.2 ± 0.6 ms after illumination *versus* 18.0 ± 0.6 ms; n= 7 rats) and that illumination alone at 365 nm has no impact on ECG parameters (RR interval: 143.4 ± 2.5 ms after illumination *versus* 143.3 ± 5.8 ms at baseline; QRS interval: 17.3 ± 0.7 ms after illumination *versus* 17.6 ± 0.6 ms; QTc: 69.0 ± 2.2 ms after illumination *versus* 69.6 ± 2.1 ms; n= 5 rats) (**Fig. 4b-d and Supplementary Table 1**).

**Figure 4.**
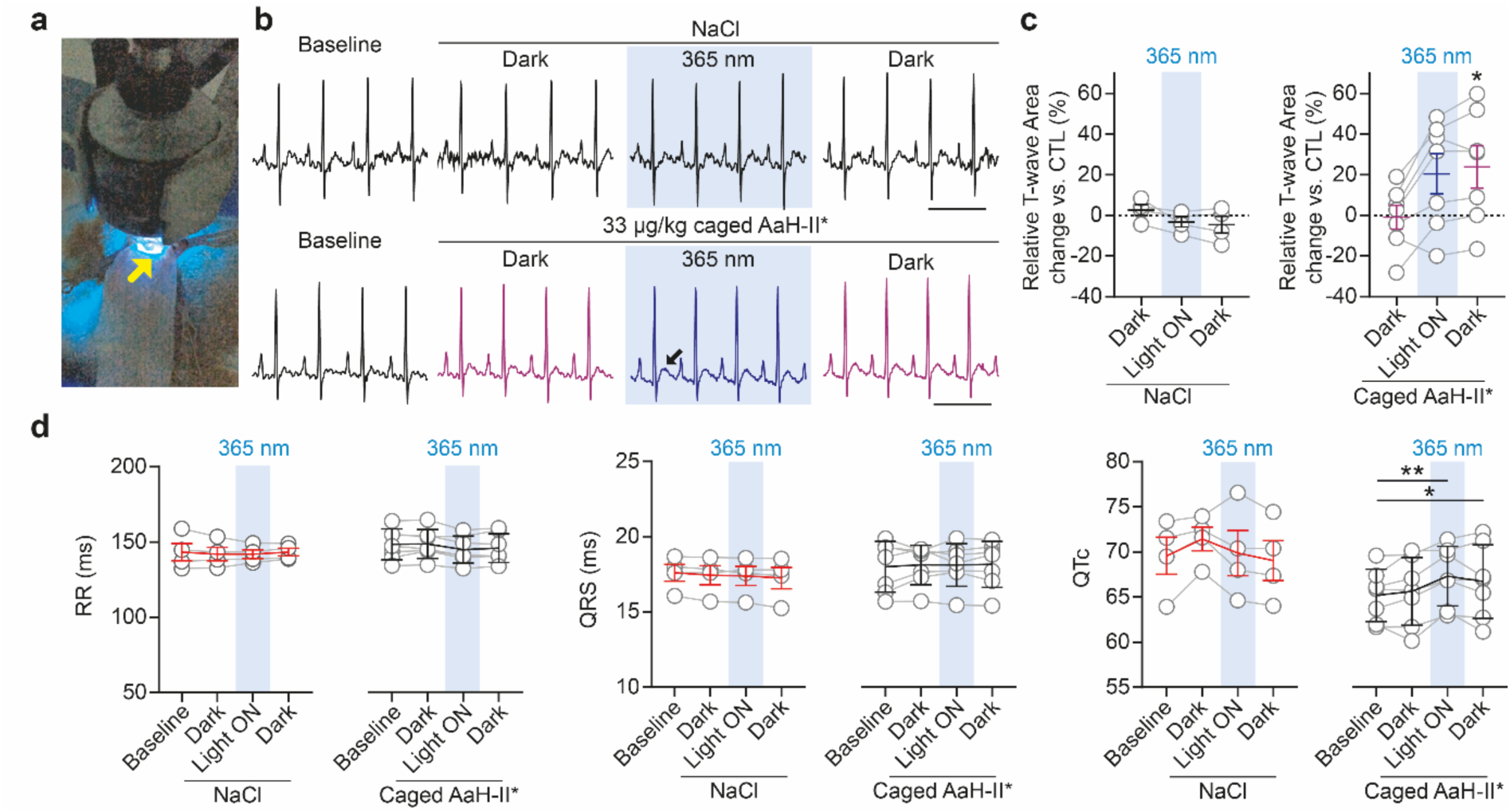
*In vivo* activation of caged AaH-II* in anesthetized rats following mini-thoracotomy. (a) Illustration of the mini-thoracotomy and external illumination of the right ventricular (RV) surface in an anesthetized rat. (b) Representative ECG traces recorded at baseline and after i.v. injection of NaCl (top panel, n = 4) or 33 µg/kg caged AaH-II* (bottom panel, n = 7) under dark conditions (3 min), after 10 min of 365 nm illumination, and 5 min post-illumination. Scale bar: 200 ms. (c) Fold change in T-wave area after NaCl injection (left panel) and T-wave area after caged AaH-II* injection (right panel) across dark, light, and post-illumination periods. (d) Quantification of RR, QRS, and QTc intervals measured under the same experimental conditions as the representative ECG traces, before and after injection in NaCl (n = 4) and caged AaH-II* (n = 7) groups. Data are presented as mean ± SEM. Statistical analysis: panel c, Friedman test followed by Dunn’s multiple-comparison test *versus* baseline using T-wave area; panel d, Friedman test followed by Dunn’s multiple-comparison test *versus* baseline. Statistical significance is indicated as *, p < 0.05 and **, p < 0.01. Source data are provided as a Source Data File.

Wireless optical device positioning and biocompatibility - To demonstrate that an implantable device can efficiently mediate photopharmacology on cardiac tissue, we used a wireless, battery-free implantable device ^8^ (**Fig. 5a**). In addition to the pacing electrode, this device contains a 405-nm µLED (Three Five Materials No. TCL20-405) with a light-emitting top surface area of 0.12 mm²and a maximal optical power of 2 mW^19^. 405 nm enables effective photoactivation of caged AaH-II* (**Supplementary Fig. 3**) and also allows better tissue penetration than 365 nm. We chose to implant the device on the upper part of the right ventricle to ensure enough light penetration of the tissue (**Fig. 5b**). We first had to validate the new surgical approach by right mini-thoracotomy (**Supplementary Fig. 5** and **6a**). The electrodes have been attached to the epicardial surface of the right ventricular outflow tract and the device receiver has been inserted in the subcutaneous space (**Fig. 5b** and **Supplementary Fig. 6**). Surgery duration was longer for implanted rats compared to sham rats (101.2 ± 6.0 min *versus* 68.8 ± 6.2 min, n = 18 and 12 rats, respectively, p < 0.01) but the post-anesthetic recovery duration was not different (15.6 ± 0.8 min *versus* 13.3 ± 1.0 min, n = 18 and 12 rats, respectively) indicating that device implantation did not affect recovery (**Supplementary Fig. 6b**). Among the implanted animals, 20 out of 23 survived the procedure. Two animals died during device implantation due to ventricular or atrial perforation, and two additional animals died within 24 hours following surgery likely due to pneumothorax induced during chest closure. Only 1 out of 13 animals died at the end of the sham procedure due to pneumothorax (**Supplementary Fig. 6c**). Post-operative follow-up (**Fig. 5c**) did not reveal any relevant alteration of the ECG parameters (**Fig. 5d-e** and **Supplementary Table 2**). We next performed Epoc analysis to assess respiratory and metabolic status 4 hours after surgery. Only a slight but significant decrease in venous sO_2_ has been observed in implanted group (87.1 ± 2.0 % after surgery *versus* 92.4 ± 1.1 % before surgery, p < 0.05, n = 10 rats) (**Fig. 5f**).

**Figure 5.**
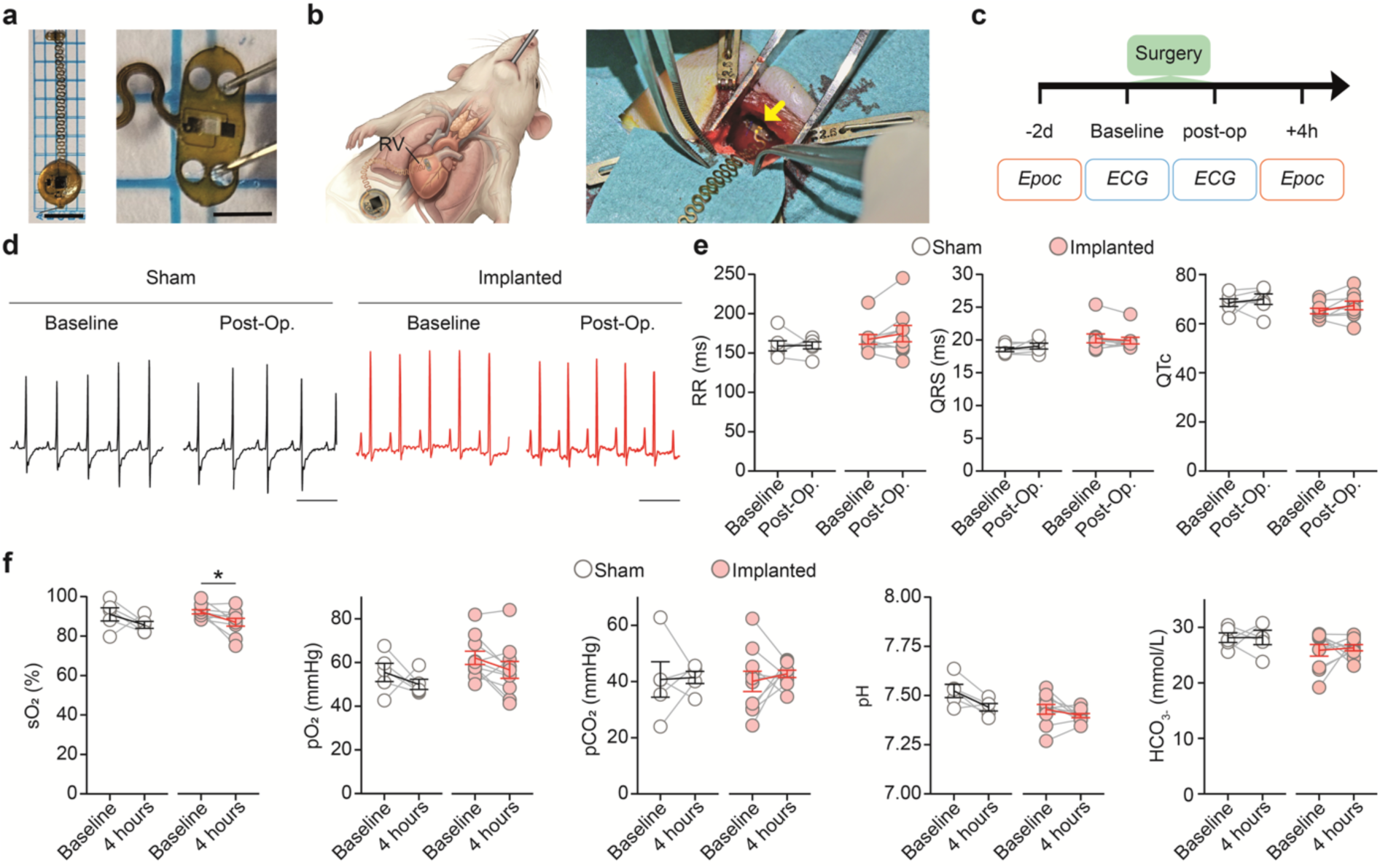
Device implantation in 10–12-week-old adult rats. (a) Images of the implantable device as a whole (left panel) and a zoomed view of the stimulation pad containing the micro-LED at its center (right panel). Scale bars = 1 cm (whole device) and 2 mm (zoom). (b) Schematic representation (left) and photograph (right panel) of the device implanted on the upper part of the right ventricular (RV) surface in intubated rats. (c) Timeline of direct post-operative follow-up including ECG and Epoc blood gas measurements. (d) Representative ECG traces recorded before and after surgery in sham (black) and implanted rats (red). Scale bar = 200 ms. (e) Quantification of RR, QRS and QTc intervals measured before and after surgery in sham (n = 6) and implanted rats (n = 5). (f) Blood gas parameters (sO₂, pO₂, pCO₂, pH, HCO₃⁻) measured before and 4 hrs after surgery in sham (n = 5) and implanted rats (n = 10). Data are presented as mean ± SEM. Statistical analysis: panels d and e, Wilcoxon matched-pairs signed-rank test. Statistical significance is indicated as *, p < 0.05 and **, p < 0.01. Source data are provided as a Source Data File.

**Figure 6.**
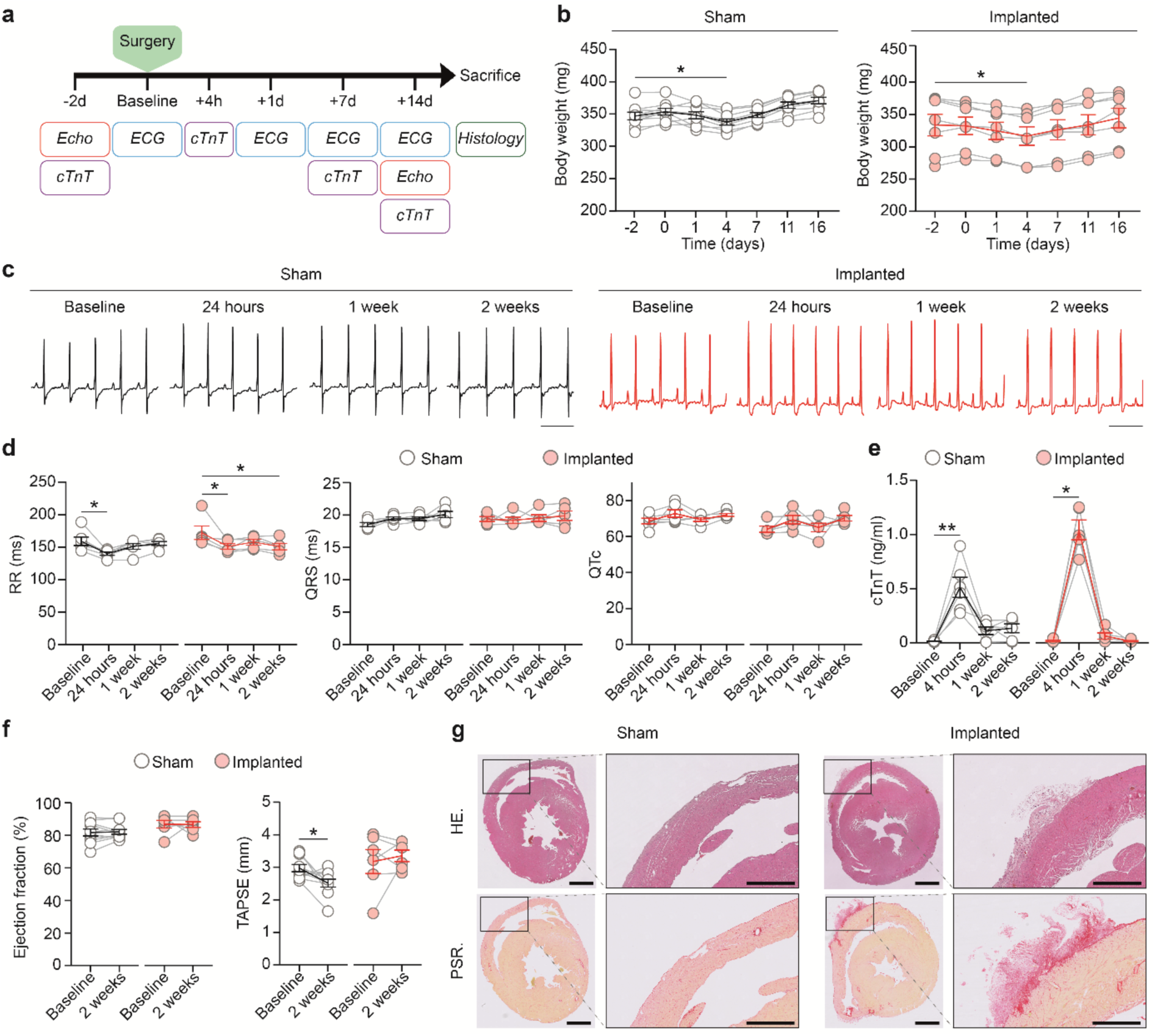
Two-week follow-up of sham and implanted rats. (a) Timeline of peri-and post-operative follow-up including echocardiography, ECG, plasma cardiac troponin T measurements, and histological assessments over 2 weeks. (b) Body weight follow-up of sham (black, n = 9) and implanted rats (red, n = 7). (c) Representative ECG traces recorded before surgery and at 24 hrs, 1 week and 2 weeks post-surgery in sham (black) and implanted rats (red). Scale bar = 200 ms. (d) Quantification of RR, QRS, and QTc intervals measured before surgery and at 24 hrs, 1 week and 2 weeks post-surgery in sham (n = 6) and implanted rats (n = 5). (e) cTnT levels measured before surgery and at 4 hrs, 1 week and 2 weeks post-surgery in sham (n = 6) and implanted rats (n = 5). (f) Quantification of left ventricular ejection fraction (left panel) and tricuspid annular plane systolic excursion (TAPSE, right panel) measured before and 2 weeks after surgery in sham (n = 10) and implanted rats (n = 6). (g) Representative heart sections collected 4 weeks post-surgery from sham rats (left panel) and implanted rats (right panel), stained with hematoxylin–eosin (top panel) and picrosirius red (bottom panel). Scale bars: 2 mm (overview) and 500 µm (RV wall). Data are presented as mean ± SEM. Statistical analysis: panels b, d and e, Friedman test followed by Dunn’s multiple-comparison test *versus* baseline; panel f, Wilcoxon matched-pairs signed-rank test. Statistical significance is indicated as *, p < 0.05, **, p < 0.01 and ***, p < 0.001. Source data are provided as a Source Data File.

We next performed a 2-week follow-up of a sub-group of both sham and implanted animals (**Fig. 6a**). Both sham and implanted rats had similar weight loss between 1 and 4 days post-surgery with a maximum weight loss of 4.7 % for sham rats and 4.9 % for implanted rats. All animals regained their initial weight 7 days post-surgery (**Fig. 6b**). We noticed slight tachycardia in sham and implanted rats 24 hours after surgery as observed by a reduction in the RR interval (Sham:158.9 ± 6.5 ms at baseline *versus* 139.9 ± 2.5 ms 24 hrs after the surgery; Implanted: 165.4 ± 10.9 ms at baseline *versus* 151.6 ± 3.8 ms 24 hrs after the surgery, p < 0.05 both, n = 6 and 5 rats, respectively) which remain present after 2 weeks only in implanted rats (RR interval:150.5 ± 4.1 ms). Other ECG parameters were not affected (**Fig. 6c-d** and **Supplementary Table 3**). To next assess cardiac injury, we evaluated myocardial injury by measuring cTnT in blood samples and observed a significant increase of cTnT levels 4 hours after surgery in both groups (14.3 ± 3.5 pg/mL at baseline versus 512.7 ± 93.11 pg/mL 4 hrs after surgery in sham rats, p<0.01, n = 5; and 14.9 ± 6.2 pg/mL at baseline *versus* 947.4 ± 83.7 pg/mL 4 hrs after surgery in implanted animals, p<0.001, n = 6). cTnT levels returned to normal levels 1-week post-surgery (0.110 ± 0.034 and 0.057 ± 0.026 ng/mL in sham and implanted rats, respectively) (**Fig. 6e**). We next evaluated cardiac function of the left and right ventricles by echocardiography. Left ventricular ejection fraction was not affected in both groups and TAPSE slightly decreased in sham group only (**Fig. 6f** and **Supplementary Table 4**). Hematoxylin-eosin and picrosirius red staining did not reveal alteration of the right myocardium despite the presence of conjunctive tissue on the epicardial side of the ventricle in the implanted area (**Fig. 6g**).

*In vivo* device-mediated modulation of cardiac activity - We next evaluated the efficacy of the device-mediated photoactivation of caged AaH-II* by using the commercially available Neurolux setup (**Fig. 7a** and **Supplementary Fig.** 7a). Before injecting the caged AaH-II* peptide, we evaluated the device integrity *in vivo* by performing an electrical 10 Hz stimulation (**Supplementary Fig. 7b**) and evaluated the effects of illumination (10 Hz, 20% duty cycle and an irradiance power of 10 mW/mm^2^) (**Fig. 7b**). Illumination alone had no effect on cardiac activity (**Fig. 7c-d** and **Supplementary Table 5**). We next injected 67 µg/kg of caged AaH-II* intravenously and did not observe any effect of this dose on ECG parameters in absence of illumination (**Supplementary Table 6**). In contrast, device-mediated photolysis of caged AaH-II* induced a significant increase in T-wave area. The caged peptide produced a relatively small fold change in T-wave area of −12.21 ± 4.38% 3 min post-injection in the dark. In contrast, a 10-min illumination led to a T-wave area change of 35.1 ± 21.9% (p < 0.01, n = 8 rats). Interestingly, this alteration in T-wave area, which reflects a change in the repolarization gradient between the illuminated and non-illuminated regions, persisted for 10 min after stopping the illumination with a value of 35.1 ± 13.3% p < 0.001, n = 8 rats). This indicates that uncaged peptide remained bound for a sufficient amount of time on its sodium channel targets in the cardiac tissue (**Fig. 7e-f** and **Supplementary Table 6**). Similar observations were made both in *ex vivo* and in open-chest experiments. We did not observe any premature ventricular contractions (PVCs) neither before or after caged AaH-II* photolysis. We speculate that this is due to the size of the illuminated cardiac region by the µLED that is too restricted to trigger PVCs. However, the absence of arrhythmias indicates that the photoactivation is regionally controlled and that the uncaged and active peptide does not diffuse away from the illuminated area in a concentration sufficient to affect the whole heart. Interestingly, a follow-up study performed in a subgroup of 3 rats showed that all rats recovered (3/3) from the anesthesia, survived 48 hrs post-procedure and displayed normal ECG parameters and no sign of AaH-II*-mediated toxicity (**Fig. 7g**). This result indicates that we were able to release enough active AaH-II* to alter locally cardiac electrical activity without inducing lethal side-effects through systemic diffusion. Paralytic and lethal effects are indeed to be expected *in vivo* if uncaged AaH-II* binds onto presynaptic Na_v_1.6 channels, as previously reported ^23^. Altogether these data demonstrate, for the first time, *in vivo* optoelectronic device-mediated modulation of cardiac activity by peptide photo-uncaging with no associated mortality.

**Figure 7.**
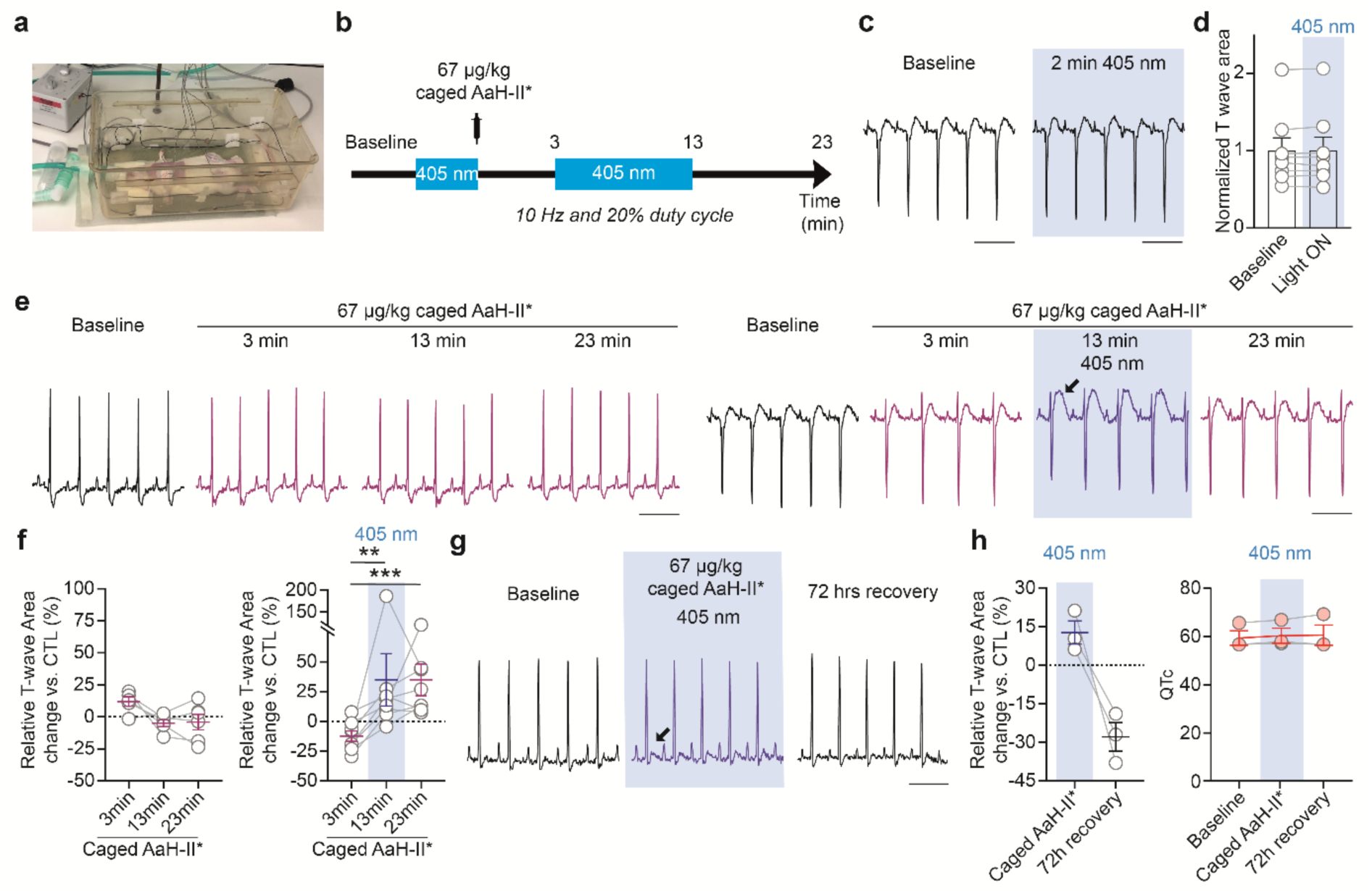
*In vivo* photoactivation of caged AaH-II* in rats using an implanted device. (a) Image of the experimental setup including the device activation system, ECG recording, gaseous anesthesia, and body temperature monitoring. (b) Experimental protocol for device-mediated peptide photoactivation. (c) Representative ECG traces recorded at baseline and after 2 min of 405 nm illumination. Scale bar = 200 ms. (d) Quantification of normalized T-wave area at baseline and during 365 nm illumination (n = 8). (e) Representative ECG traces recorded at baseline and at 3, 13 and 23 min after i.v. injection of 67 µg/kg caged AaH-II* (left panel). Representative ECG traces recorded at baseline, 3 min after i.v. injection of 67 µg/kg caged AaH-II*, after 10 min of illumination and after 10 min in the dark (right panel). Scale bar = 200 ms. (f) Relative T-wave area fold change normalized to baseline under non-illuminated conditions (baseline and 3, 13, and 23 min after i.v. injection of 67 µg/kg caged AaH-II*; left panel, n = 5) and illuminated conditions (baseline, 3 min after i.v. injection of 67 µg/kg caged AaH-II*, after 10 min of illumination, and after 10 min in the dark; right panel, n = 8). (g) Representative ECG traces recorded during control conditions, after 10 min of caged AaH-II* photoactivation, and after 72 hrs recovery. Scale bar = 200 ms. (h) Relative T-wave area fold change normalized to baseline (left panel, n = 3) and QTc quantification at baseline, after 10 min of caged AaH-II* photoactivation, and after 72 hrs recovery (right panel, n = 3). Data are presented as mean ± SEM. Statistical analysis: panel d, Wilcoxon matched-pairs signed-rank test; panel f, Friedman test followed by Dunn’s multiple-comparison test using T-wave area; panel h, Friedman test followed by Dunn’s multiple-comparison test versus baseline. Statistical significance is indicated as *, p < 0.05, **, p < 0.01 and ***, p < 0.001. Source data are provided as a Source Data File.

## Discussion

This study provides the first demonstration of *in vivo* cardiac photopharmacology using a caged peptide driven by an implantable wireless device, establishing a new paradigm in which drug activity is controlled in time and space after systemic delivery. Pharmacological activity is thereby uncoupled from systemic exposure and restricted to a defined myocardial region. Building on our previous demonstration of region-selective photopharmacology in the right atrium ^22^, we now extend this control to the right ventricle *in vivo* and with an optoelectronic device. At the organ scale, photoactivation induces localized alterations of repolarization without affecting conduction or heart rate. The effect is therefore spatially confined and mechanistically consistent with selective modulation of sodium-channel inactivation. *In vivo*, intravenous caged peptide remains inert, whereas targeted illumination, either fiber-based or delivered *via* an implanted wireless device, elicits controlled changes in T-wave morphology and QTc without systemic electrophysiological disturbance. The absence of effects with light alone further validates the specificity and safety of the approach. Despite the invasiveness of implantation, rats showed rapid recovery and preserved cardiac function supporting the biocompatibility of the system and showing that flexible, wireless, fully implantable optoelectronic interfaces can be tolerated chronically on the rodent epicardium in line with previous reports ^6–8,32^. The absence of device-mediated arrhythmias highlights the importance of spatial scale in shaping electrophysiological outcomes, consistent with the well-established notion that a critical mass of electrically perturbed myocardium is required to sustain reentry, whereas sub-organ activation predominantly generates benign repolarization gradients ^33^.

These features position peptide photoactivation as a powerful alternative to conventional pharmacology and optogenetics, particularly for applications requiring non-genetic, reversible, and region-specific interventions as previously illustrated for smaller compounds active onto muscarinic receptors ^34^, beta-adrenergic receptors ^35^ or Na_v_1.5 ^36^. Beyond proof-of-concept, this work defines a scalable framework for precision modulation of excitable tissues at sub-organ resolution. It decouples efficacy from off-target toxicity and paves the way for integration into closed-loop bioelectronic systems for adaptive therapies, in which on-board sensing and computation could gate drug photoactivation in response to real-time ECG signals ^7,37^.

Several improvements in the technology should nevertheless be considered and expected in the future. First, activation in the UV range (365-405 nm) limits tissue penetration and may restrict applications to tissues of limited thickness. The development of red-shifted caging groups is therefore a key next step, with promising scaffolds already reported in the visible-to-NIR range, including water-soluble BODIPY photocages activatable at 633 nm ^38^. Second, the illuminated myocardial volume is constrained by the current implant geometry, and further iterations of the optoelectronic devices will require application-tailored designs. Multi-site µLED arrays with subject-specific geometries ^39^ and machine-learning–guided ultrathin epicardial arrays ^7^ have already been developed for optogenetics, albeit with complex fabrication. They provide a blueprint for addressable µLED matrices delivering dose-and region-tunable uncaging patterns. Finally, long-term studies are needed to assess chronic biocompatibility and device stability over time in moving animals. Newer designs now achieve sustained optical cardiac neuromodulation for months post-myocardial-infarction in a canine model ^40^, outlining a realistic path to chronically stable cardiac photopharmacology implants.

This study lays the groundwork for next-generation precision therapeutics that integrate chemistry and bioelectronics to control cardiac electrical activity with unprecedented spatial and temporal resolution.

## Methods

### Ethical Statement

All animal care and experimental procedures were performed in an animal facility that has been accredited by the French Ministry of Agriculture. The experimental procedures were approved by the regional ethic committee (#38691-202209261659333 v4 CEEA-006 Pays de la Loire, France) according to the Directive 2010/63/EU of the European Union.

### Molecular modelling

The impact of adding the Nvoc group at lysine 62 of AaH-II-R^62^K (termed AaH-II*) on the interactions between the peptide and the Na_V_1.7-Na_V_PaS channel was predicted using available structures of native AaH-II (PDB code: 6NT4) ^30^. Discovery Studio (version 21.1.0.20298; Dassault System) was used design caged AaH-II* (AaH-II-R^62^K-Nvoc). After applying a fast, Dreiding-like forcefield in order to clean geometry of designed peptide, steric hindrance induced by the addition of the Nvoc group to AaH-II* was analyzed with the “Show_bumps” module in PyMol (http://www.pymolwiki.org/index.php/Show_bumps). Red circles indicate steric clashes related to van der Waals interactions between peptide and sodium channel.

### Synthesis of peptides analogues

Peptides were assembled stepwise using Fmoc-based Solid Phase Peptide Synthesis (SPPS) on a PTI Symphony synthesizer at a 0.05 mmol or 0.1 mmol scale on 2-chlorotrityl chloride polystyrene resin (substitution approximately 1.6 mmol/g) by Smartox Biotechnology (Saint-Egrève, France). For the caged peptide, Fmoc-L-Lysine residues at position 62 was replaced by Fmoc-L-Lys(Nvoc)-OH (Iris Biotech, Marktredwitz, Germany) during assembly. The Fmoc protecting group was removed using 20% piperidine in DMF and free amine was coupled using tenfold excess of Fmoc amino acids and HCTU/DIEA activation in NMP/ DMF (3×15 min). The peptide was de-protected and cleaved from the resin with TFA/H_2_O/1,3-dimethoxybenzene(DMB)/TIS/2,2′-(ethylenedioxy)diethanethiol (DODT) 85.1/5/2.5/3.7/3.7 (vol.), then precipitated out in cold diethyl ether. The resulting white solids were washed twice with diethyl ether, re-suspended in H_2_O/acetonitrile, and freeze dried to afford crude linear peptide. Oxidative folding of the crude linear caged AaH-II* was successfully conducted at RT in the conditions optimized for the caged AaH-II* using a concentration of 0.1 mg/mL in 0.1 M Tris buffer at pH 8.0 containing 10% of DMSO. Purification of the peptide was performed on Chromolith^®^ High Resolution RP18e column (100 ×4.6 mm) using a 5-65% buffer B run in 14 min at 2 mL/min. Solvent system: A: H_2_O, 0.1%TFA and B: MeCN 0.1%TFA.

### LC-ESI-MS analyses

Liquid chromatography–electrospray ionization mass spectrometry (LC-ESI-MS) analyses were performed using an Agilent 1290 UPLC system coupled to an Accurate Mass QTOF 6530 mass spectrometer (Agilent Technologies). When chromatographic separation was required, peptide samples (5 µL, approximately 10 µg/mL) were injected onto an Ascentis Express C18 column (90 Å, 2.0 µm particle size, 2.1 × 100 mm; Supelco) maintained at 70°C. Elution was carried out at a flow rate of 0.5 mL/min using a linear gradient from 2% to 70% solvent B over 11 min, where solvent A consisted of H₂O/formic acid (99.9:0.1, v/v) and solvent B of acetonitrile/formic acid (99.9:0.1, v/v). For samples not requiring chromatographic separation, direct infusion was performed at a flow rate of 0.6 mL/min using an H₂O/acetonitrile solution supplemented with 0.1% formic acid. Mass spectra were acquired in positive ionization mode using a Dual ESI source over an m/z range of 100–1700 and processed with Agilent MassHunter software (version 10.0). Instrument settings were as follows: capillary voltage, 4.0 kV; fragmentor voltage, 100 V; gas temperature, 350°C; drying gas flow, 12 L/min; and nebulizer pressure, 50 psig.

### Analytical RP-HPLC analyses

Analytical RP-HPLC was performed using an SPD M20-A system (Shimadzu) with a Luna OmegaPS C18 column (4.6 ×250 mm, 5 μm, 100 Å). Twenty μL (corresponding to 7 μg of material) was loaded and a 5-60% acetonitrile gradient (0.1% TFA v/v) was applied over 35 min at room temperature to detect analytes by UV absorbance at 214 nm. Illumination of samples was performed at 365 at 41.8 mW/cm^2^ for 30 sec to 30 min using a CoolLED pE4000 light source (CoolLED, UK). Flash energy has been measured using a highly sensitive thermal power head (S401C, ThorLabs).

### Cell cultures

HEK293 cells stably expressing the human (h)Na_v_1.2 (XM_017004655) channels were cultured in Dulbecco’s Modified Eagle’s Medium (DMEM) containing 1 mM sodium pyruvate and 4.5 g/L glucose, supplemented with 2 mM glutamine, 10% fetal bovine serum (FBS), 10 U/mL penicillin, and 10 μg/mL streptomycin (Gibco, USA). Selection pressure was maintained with 800 μg/mL G418 (Roche, Switzerland). Cultures were incubated at 37°C under 5% CO₂ and maintained at 60-80% confluence. For patch-clamp recordings, cells were detached with trypsin and floating single cells were diluted (∼300,000 cells/mL) in medium contained (in mM): 140 mM, 4 KCl, 2 CaCl_2_, 1 MgCl_2_, 5 glucose and 10 HEPES (pH 7.4, osmolarity 298 mOsm).

### High-throughput patch-clamp recordings and pharmacological assays

Whole-cell recordings were used to investigate the effects of caged AaH-II* on HEK293 cells expressing hNa_v_1.2. Automated patch-clamp recordings were performed using the SyncroPatch 384PE from Nanion (München, Germany). Chips with single-hole medium resistance of 4.37 ± 0.16 MΩ (n = 384) were used for recordings. Pulse generation and data collection were performed with the PatchControl384 v1.5.2 software (Nanion) and the Biomek v1.0 interface (Beckman Coulter). Whole-cell recordings were conducted according to the recommended procedures of Nanion. Prior to each experiment, cells were isolated as previously described. Cells were stored in a cell hotel reservoir at 10°C with shaking speed at 200 RPM. After initiating the experiment, cell catching, sealing, whole-cell formation, liquid application, recording, and data acquisition were all performed sequentially and automatically. Intracellular solution contained (in mM): 10 CsCl, 110 CsF, 10 NaCl, 10 EGTA, and 10 HEPES (pH 7.2, osmolarity 280 mOsm). Extracellular solution was (in mM): 140 NaCl, 4 KCl, 2 CaCl_2_, 1 MgCl_2_, 5 glucose and 10 HEPES (pH 7.4, osmolarity 298 mOsm). Recordings were performed at room temperature (18-22°C). The holding potential was set at-100 mV, and the sampling rate was 20 kHz. Compounds were tested at +10 mV with 50 ms pulse duration repeated every 5 seconds. The peptide was diluted in extracellular solution containing 0.3% bovine serum albumin (BSA). Effects of peptide on current properties were measured at equilibrium after 15 minutes of peptide application time. In order to evaluate the ability of photoactivation to restore AaH-II* effects on hNa_v_1.2, after 2 min of control condition, caged AaH-II* was added to the external buffer and effects were recorded for 100 sec prior to a 250 sec-duration illumination at 45 mW/cm^2^ for photocleavage induction. Acquired data were exported from DataControl384 (Version 1.8.0; Nanion Technologies, Germany) and analysis were performed using custom R scripts in RStudio (Version 1.4.1717). Concentration-response curves were plotted and fitted using GraphPad Prism (Version 8.4.2), providing EC₅₀ values of caged AaH-II* and uncaged AaH-II* for hNa_v_1.2. Because AaH-II* slows sodium channel inactivation and thereby enhances the late sodium current, increasing the total Na⁺ influx during depolarization, we quantified Na⁺ entry by calculating the area under the curve (AUC), which reflects the total charge carried by Na⁺ during the depolarizing step.

### Ex vivo photoactivation on Langendorff-perfused hearts

Eleven-week-old male Wistar-Han rats were placed in an induction chamber and anesthetized with 5% isoflurane delivered at 1.2 L/min of air. Euthanasia was then performed via intraperitoneal injection of a lethal dose of pentobarbital (140 mg/kg; Euthasol®) combined with sodium heparin (2,500 IU). Following euthanasia, hearts were rapidly excised and rinsed in a modified cold (4°C) Krebs-Henseleit (KH) solution containing (in mM): 116 NaCl, 5 KCl, 1.1 MgSO₄·7H₂O, 0.35 NaH₂PO₄, 27 NaHCO₃, 10 glucose. Hearts were quickly cannulated and perfused in a retrograde fashion through an aortic cannula with warm (37-38°C) oxygenated (95% O₂ / 5% CO₂) modified KH solution supplemented with 1.8 mM CaCl₂, 0.2 mM pyruvate and 1 mM lactate, at a constant flow rate of 12 mL/min. Cardiac surface electrical activity was measured using a MultiElectrode Array (MEA) positioned over the right ventricular outflow tract. Signals were recorded using a 64-electrode array (0.2-mm electrode diameter, 0.36-mm interelectrode spacing) connected to an EMS64-USB-1003 amplifier (MappingLab, Oxford, UK). For each electrode, ventricular electrical signal was characterized by a prominent depolarization wave followed by a smaller repolarization wave. Simultaneously, single-lead ECG recordings were acquired to monitor global cardiac electrical activity. Signals were sampled at 10 kHz per channel and acquired using EMapRecord5 software (v5.8.61, MappingLab, Oxford, UK). After a 10-min stabilization and control period of KH perfusion, 10 nM caged AaH-II* was perfused for 13 min. Illumination at 365 nm at 75 mW/cm^2^ was applied at 3 min of perfusion using a CoolLED pE4000 light source (CoolLED, UK) for 10 min. All experiments were conducted in the dark to prevent unwanted uncaging. Data were analyzed using EMapScope5 software (v5.9.43, MappingLab, Oxford, UK). Thirty-second recording segments were analyzed at the end of the control period, after 3, 6 and 9 minutes of perfusion with caged AaH-II*. Effects of the compound were evaluated by assessing RR intervals between consecutive signals, conduction velocity, repolarization wave duration, and repolarization wave area (analyzed in raw form and expressed as the mean relative fold change from baseline).

### In-vivo open thorax photoactivation

Ten-to twelve-week-old male rats were anesthetized with sevoflurane (8% sevoflurane, FiO_2_ = 0.21, 2 L/min for induction and 4% sevoflurane, FiO_2_ = 0.21, 0.6 L/min for maintenance). Animals were intubated using a custom-made 14G metallic probe and mechanically ventilated throughout the procedure (tidal volume 8-9 mL/kg, 55 ± 4 cycles/min; Minivent Type 845, Hugo Sachs Electronik). Body temperature was monitored using a rectal probe and maintained at 37.0 ± 1.0°C using a heating pad. Continuous ECG monitoring was performed using subcutaneous limb electrodes connected to a digital acquisition system (IOX, EMKA Technologies, Paris, France). To expose the heart, a right-sided thoracotomy was performed to expose the heart RV surface. The right ventricular (RV) surface of the heart was exposed by performing a right-sided thoracotomy. The ribs were maintained apart using a surgical retractor. Following a stabilization period, animals received an intravenous injection through the dorsal penile vein of either saline (NaCl) or caged AaH-II* (33 µg/kg in 120 µL). Three minutes after injection, the exposed heart was illuminated for 10 min using a 365 nm light source (75 mW/cm²) delivered externally above the heart through an optical fiber coupled to a CoolLED pE4000 illumination system (CoolLED, UK), identical to that used for isolated heart experiments. Illumination was applied directly onto the cardiac surface without contact with the tissue. ECG recordings were continuously acquired before, during, and for 5 min following termination of illumination. At the end of the protocol, animals were euthanized using i.v. injection of a lethal dose of pentobarbital.

### Surgical procedure

Surgeries and post-operative care were conducted in accordance with good surgical and aseptic practice, under sterile drapes, in a clean area, using sterilized equipment. Animals were randomly assigned to sham (no implant) or implanted groups. Ten-to twelve-week-old male rats were premedicated 20 min before the surgery with buprenorphine (0.05 mg/kg s.c.) and meloxicam (2 mg/kg s.c.). Ophthalmic gel was applied (Ocry-gel, TMV) and 5 mL/kg of 0.9% NaCl solution was injected subcutaneously to prevent dehydration. The animals were intubated using a custom-made 14G metallic probe under general anesthesia using sevoflurane (8% sevoflurane, FiO_2_ = 0.21, 2 L/min for induction and 4% sevoflurane, FiO_2_ = 0.21, 0.6 L/min for maintenance), transferred to the surgical field and placed in dorsal recumbency. Animals were mechanically ventilated during the procedure (tidal volume of 8 to 9 mL/kg and 55 ± 4 cycles per min; Minivent Type 845, Hugo Sachs Electronik), allowing the maintenance of the respiratory function and the anesthesia. Six-lead ECG monitoring was performed with 25-gauge subcutaneous electrodes on the limbs coupled to a digital amplifier and recording system (IOX 1.585; EMKA Technologies, Paris, France). Body temperature was monitored and maintained at 37.0 ± 1.0°C using a heating pad and a rectal temperature probe. To access the right ventricle, right-side mini-thoracotomy was performed in the 3^rd^ intercostal space **(Supplementary figure 6)**. Pectoral muscles were separated and pulled apart, exposing the ribs. A Heiss retractor was used to create an operative window providing access to the heart. To expose the right ventricle, the right pulmonary lobes were gently displaced laterally, and the pericardium was carefully opened using surgical forceps. The device was then sutured on the upper part of the right ventricle using 7-0 Prolene suture as previously described ^41^. Thoracotomy was closed using a 4-0 Prolene suture, while recreating the pleural vacuum to ensure proper lung function. The receptor portion of the device was inserted beneath the skin of the abdomen and ventral skin was close using 5-0 Prolen suture. Finally, the animals received injection of 0.9% NaCl (5 mL/kg s.c.) and Buprenorphine (0.05 mg/kg s.c.) and were extubated upon recovery after the termination of anesthesia. Animals recovered in an infant incubator (temperature adjusted to 28°C) for 2 days while receiving post-operative follow-up. Buprenorphine (0.05 mg/kg s.c.) and meloxicam (2 mg/kg s.c.) were injected 4 hrs, 24 and 48 hrs after the surgery, respectively. The device functionality was tested 4 hrs after the surgery by triggering electrical stimulations (pulses of 20 ms at 10 Hz for 30 sec). Implanted and non-implanted sham animals weighed daily for up to 2 weeks following implantation.

### Cardiac follow-up of animals

Sham and implanted rats were monitored for recovery over 2 weeks. Blood gas parameters (pO_2_, pCO_2_, sO_2_, pH) were analyzed using Epoc blood analysis device (epoc^®^ 3.40.3, Siemens Heathineers, Germany) and BGEM BUN cards by collecting 100 µL of venous blood from retro-orbital sinus or subclavian veins in anesthetized rats 24 to 48 hrs before the surgery (baseline) and 24 hrs after the surgery. ECG was performed as previously described 24 hrs, 1 week and 2 weeks after the surgery. PR, QRS, QT, and RR intervals were measured using ECGAuto software (ECGAuto v3.5.5.28, EMKA Technologies). The end of the T wave was defined as the point at which signal returned to the isoelectric line. QT intervals were corrected for heart rate using the formula (Bazett’s formula normalized to average rat RR of 150 ms): QTc=QT/(RR/150)^1/2^ with QT and RR measured in ms ^42^. Transthoracic echocardiographic studies were performed under general anesthesia using sevoflurane 2 weeks after surgery and compared to recordings performed 48 hrs prior surgery (Vevo 3100, with the MX201 transducer, 10-22MHz, FUJIFILM VisualSonics, Canada), following current recommendations^43^. M-mode was used in parasternal short and long axis to measure thickness of the left ventricle and to estimate the left ventricular end-diastolic volume (EDV) and end-systolic volume (ESV). The left ventricular ejection fraction was subsequently calculated using EDV and ESV: (EDV-ESV)/EDV*100. The right ventricle function was determined by measuring Tricuspid Annular Plane Systolic Excursion (TAPSE). All parameters were analyzed using Vevo Lab analysis software 5.10 (FUJIFILM VisualSonics, Canada). Cardiac TnT (cTnT) level was measured from blood plasma collected from retro-orbital sinus or subclavian veins 24 to 48 hrs before the surgery (baseline) and 4 hrs, 1 week and 2 weeks after the surgery. Blood samples collected in heparinized aliquots were centrifugated during 10 min at 10,000 rpm to separate plasma from other blood component and cTnT levels were measured by electrochemiluminescence immunoassay (ECLIA) technique using a Cobas e 801 (Roche, France).

### Histological analysis

Sham and implanted rat hearts were excised 4 weeks after the surgery, fixed in 10% formalin and embedded in paraffin. Coronal hearts sections of 7-µm thickness (HLM 355S, Microm Microtech France) were stained with Hematoxylin-Eosin and picrosirius red as previously described ^44^.

### In vivo photoactivation of caged peptide

Implanted rats were placed in a custom-made experimental cage to allow simultaneous ECG recording and device activation while maintaining animal anesthesia using sevoflurane and monitoring body temperature. Rats’ anesthesia and ECG recordings were performed as previously described ^22^. The functionality of the device was verified prior to each experiment by triggering electrical stimulations (10 Hz with 5% duty cycle for 30 sec) and next, a 2-min baseline ECG was recorded prior to intravenous injection of 67 µg/kg (120 µL) caged AaH-II* in dorsal penile vein. Three min after injection, the device µLED was activated (10 Hz with 20% duty cycle) during 10 min and ECG recordings continued during the ensuing 10 min following termination of peptide activation. ECGs were analyzed as previously described. Additionally, area of T wave is measured between S wave and the end of T wave to monitor AaH-II*-mediated effects. Individual ECG data were averaged over 10 sec for each period (baseline, 3, 13 and 23 min after peptide injection) to reduce variability of the measure.

### Statistics and data analyses

Values are represented as mean ± SEM. Statistical analyses were performed using GraphPad Prism version 8.4.1 (GraphPad Software, San Diego, CA, USA). As several datasets did not meet normality assumptions, non-parametric tests were used throughout the study. Exact sample sizes (n) are indicated in the corresponding figure legends. A p-value lower than 0.05 was considered significant. In patch-clamp experiments, the Friedman test followed by Dunn’s multiple-comparisons test was used to assess differences in peptide potency for hNa_v_1.2 at 1 nM. The Mann–Whitney U test was used to compare the dynamic photoactivation of caged AaH-II* on sodium currents. All *ex vivo* MEA-mediated and *in vivo* open-thorax ECG parameter analyses were performed using the Friedman test followed by Dunn’s multiple-comparisons post hoc test for repeated measures. Post-operative parameters, including ECG and EPOC measurements, were analyzed using the Mann–Whitney U test. For the 2-week follow-up study, parameters including four or more time points, such as body weight, ECG parameters, and cTnT levels, were analyzed using the Friedman test followed by Dunn’s multiple-comparisons test versus baseline. Echocardiographic parameters were analyzed using the Wilcoxon matched-pairs signed-rank test. Differences between sham and implanted groups were additionally assessed using the Mann-Whitney U test. Effects of device-mediated illumination on cardiac activity were assessed using the Wilcoxon matched-pairs signed-rank test. The significance of peptide photoactivation was evaluated using the Friedman test followed by Dunn’s multiple-comparisons post hoc test performed within each column on raw data values. Recovery 72 hrs after photoactivation was analyzed using the Friedman test followed by Dunn’s multiple-comparisons test versus baseline.

## Data availability

The data that support the findings of this study and analytical tools are available here: https://github.com/jerome-montnach/AaHII-implant.

## Acknowledgements

We thank the animal facility care unit (UTE IRS-UN) for animal housing and management. The authors also thank the CRNH-Ouest Mass Spectrometry Core Facility (l’institut du thorax, Nantes, France), AMaCC (CEISAM, Nantes, France) and Neurolux company for their technical support. We acknowledge THERASSAY core facility (SFR Bonamy, l’institut du thorax INSERM UMR 1087, CNRS UMR 6291, Nantes, France) member of the Scientific Interest Group (GIS) Biogenouest and IBISA, for the use of the echograph and their technical support.

## Funding

J. Montnach thanks the Agence Nationale de la Recherche (ANR) for its financial support (WOCAR grant N° ANR-23-CE19-008-C1), including the fellowship of M. Derelli. This work was also supported by the PULSAR program from Région Pays de la Loire, by the “International Emerging Action” program from CNRS, by the Fondation Leducq in the frame of its program of ERPT equipment support (to MDW to purchase an automated patch-clamp system and by a European FEDER grant in support of the automated patch-clamp infrastructure).

## Author contributions statement

M.D. and H.M. performed cell culture, automated patch-clamp, RP-HPLC, ex vivo and in vivo experiments and analyses. A.H and A.P. performed all surgical experiments. M.C.B performed cell culture and patch-clamp experiments and analysis. V.A and P.A. performed echocardiography experiments and analyses. C.Z. and R.B. performed the chemical synthesis of peptides. E.B. performed cTnT dosage. M.D and J.M made the figures. J.M and M.D.W conceived and managed the project. J.M wrote the original draft of the manuscript. P.G., F.C., M.D.W and J.M reviewed and edited the manuscript.

## Notes

### Competing Interest Statement

The authors have declared no competing interest.

## References

1 Merino, J. L. et al. Practical compendium of antiarrhythmic drugs: a clinical consensus statement of the European Heart Rhythm Association of the European Society of Cardiology. Europace: European pacing, arrhythmias, and cardiac electrophysiology: journal of the working groups on cardiac pacing, arrhythmias, and cardiac cellular electrophysiology of the European Society of Cardiology 27 (2025). 10.1093/europace/euaf076

2 Nanasi, P. P., Pueyo, E. & Virag, L. Editorial: Perspectives of Antiarrhythmic Drug Therapy: Disappointing Past, Current Efforts, and Faint Hopes. Front Pharmacol 11, 1116 (2020). 10.3389/fphar.2020.01116

3 Sorodoc, V. et al. Amiodarone Therapy: Updated Practical Insights. J Clin Med 13 (2024). 10.3390/jcm13206094

4 Mond, H. G. & Proclemer, A. The 11th World Survey of Cardiac Pacing and Implantable Cardioverter-Defibrillators: Calendar Year 2009-A World Society of Arrhythmia’s Project. Pacing and Clinical Electrophysiology 34, 1013–1027 (2011). 10.1111/j.1540-8159.2011.03150.x

5 Al-Khatib, S. M. et al. 2017 AHA/ACC/HRS Guideline for Management of Patients With Ventricular Arrhythmias and the Prevention of Sudden Cardiac Death. Circulation 138, e272–e391 (2018). 10.1161/CIR.0000000000000549

6 Chang, Z. et al. Fully Implantable Wireless Cardiac Pacing and Sensing System Integrated with Hydrogel Electrodes. Adv Sci (Weinh) 11, e2401982 (2024). 10.1002/advs.202401982

7 Ausra, J. et al. Wireless, fully implantable cardiac stimulation and recording with on-device computation for closed-loop pacing and defibrillation. Science advances 8, eabq7469 (2022). 10.1126/sciadv.abq7469

8 Gutruf, P. et al. Wireless, battery-free, fully implantable multimodal and multisite pacemakers for applications in small animal models. Nature communications 10, 5742 (2019). 10.1038/s41467-019-13637-w

9 Liu, Y., Wang, T. & Wang, W. Photopharmacology and photoresponsive drug delivery. Chem Soc Rev (2025). 10.1039/d5cs00125k

10 Hull, K., Morstein, J. & Trauner, D. In Vivo Photopharmacology. Chemical reviews 118, 10710–10747 (2018). 10.1021/acs.chemrev.8b00037

11 Nyns, E. C. A. et al. Optical ventricular cardioversion by local optogenetic targeting and LED implantation in a cardiomyopathic rat model. Cardiovasc Res (2021). 10.1093/cvr/cvab294

12 Bruegmann, T., Beiert, T., Vogt, C. C., Schrickel, J. W. & Sasse, P. Optogenetic termination of atrial fibrillation in mice. Cardiovasc Res 114, 713–723 (2018). 10.1093/cvr/cvx250

13 Nyns, E. C. A. et al. Optogenetic termination of ventricular arrhythmias in the whole heart: towards biological cardiac rhythm management. European heart journal 38, 2132–2136 (2017). 10.1093/eurheartj/ehw574

14 Bruegmann, T. et al. Optogenetic defibrillation terminates ventricular arrhythmia in mouse hearts and human simulations. J Clin Invest 126, 3894–3904 (2016). 10.1172/JCI88950

15 Bingen, B. O. et al. Light-induced termination of spiral wave arrhythmias by optogenetic engineering of atrial cardiomyocytes. Cardiovasc Res 104, 194–205 (2014). 10.1093/cvr/cvu179

16 Tochitsky, I., Kienzler, M. A., Isacoff, E. & Kramer, R. H. Restoring Vision to the Blind with Chemical Photoswitches. Chemical reviews 118, 10748–10773 (2018). 10.1021/acs.chemrev.7b00723

17 Casson, R. J., et al. in Nat Med (2026).

18 Mourot, A. et al. Rapid optical control of nociception with an ion-channel photoswitch. Nat Methods 9, 396–402 (2012). 10.1038/nmeth.1897

19 Kim, M. et al. Wireless, battery-free, remote photoactivation of caged-morphine for photopharmacological pain modulation without side effects. Biosens Bioelectron 281, 117440 (2025). 10.1016/j.bios.2025.117440

20 Fehrentz, T. et al. Optical control of cardiac electrophysiology by the photochromic ligand azobupivacaine 2. British journal of pharmacology (2024). 10.1111/bph.17394

21 Fehrentz, T. et al. Optical control of L-type Ca(2+) channels using a diltiazem photoswitch. Nature chemical biology 14, 764–767 (2018). 10.1038/s41589-018-0090-8

22 Montnach, J. et al. Optical Control of Cardiac Rhythm by In Vivo Photoactivation of an ERG Channel Peptide Inhibitor. Circulation research 133, 535–538 (2023). 10.1161/CIRCRESAHA.123.322880

23 Montnach, J. et al. In vivo spatiotemporal control of voltage-gated ion channels by using photoactivatable peptidic toxins. Nature communications 13, 417 (2022). 10.1038/s41467-022-27974-w

24 Laughner, J. I. et al. A fully implantable pacemaker for the mouse: from battery to wireless power. PLoS One 8, e76291 (2013). 10.1371/journal.pone.0076291

25 Gutruf, P. & Rogers, J. A. Implantable, wireless device platforms for neuroscience research. Current opinion in neurobiology 50, 42–49 (2018). 10.1016/j.conb.2017.12.007

26 Nussinovitch, U. & Gepstein, L. Optogenetics for in vivo cardiac pacing and resynchronization therapies. Nat Biotechnol 33, 750–754 (2015). 10.1038/nbt.3268

27 Ouyang, W. et al. A wireless and battery-less implant for multimodal closed-loop neuromodulation in small animals. Nat Biomed Eng 7, 1252–1269 (2023). 10.1038/s41551-023-01029-x

28 Bhatia, A. et al. Wireless Battery-free and Fully Implantable Organ Interfaces. Chemical reviews 124, 2205–2280 (2024). 10.1021/acs.chemrev.3c00425

29 Millet, H. et al. Late INa activation of cardiac TTX-sensitive sodium channels by AaH-II induces an arrhythmogenic phenotype. bioRxiv (2025). 10.1101/2025.09.17.676709

30 Clairfeuille, T. et al. Structural basis of alpha-scorpion toxin action on Nav channels. Science 363 (2019). 10.1126/science.aav8573

31 Johnson, E. C. & Kent, S. B. Synthesis, stability and optimized photolytic cleavage of 4-methoxy-2-nitrobenzyl backbone-protected peptides. Chemical communications, 1557– 1559 (2006). 10.1039/b600304d

32 Choi, Y. S. et al. Fully implantable and bioresorbable cardiac pacemakers without leads or batteries. Nat Biotechnol (2021). 10.1038/s41587-021-00948-x

33 Nyns, E. C. A. et al. An automated hybrid bioelectronic system for autogenous restoration of sinus rhythm in atrial fibrillation. Sci Transl Med 11 (2019). 10.1126/scitranslmed.aau6447

34 Riefolo, F. et al. Optical Control of Cardiac Function with a Photoswitchable Muscarinic Agonist. Journal of the American Chemical Society 141, 7628–7636 (2019). 10.1021/jacs.9b03505

35 Duran-Corbera, A. et al. A photoswitchable ligand targeting β1-adrenoceptor enables light-control of the cardiac rhythm. (2022). 10.1101/2022.03.06.483174

36 Liu, S. et al. in Nature communications Vol. 17 (2026).

37 Choi, Y. S. et al. A transient, closed-loop network of wireless, body-integrated devices for autonomous electrotherapy. Science 376, 1006–1012 (2022). 10.1126/science.abm1703

38 Poryvai, A., Galkin, M., Shvadchak, V. & Slanina, T. Red-Shifted Water-Soluble BODIPY Photocages for Visualisation and Controllable Cellular Delivery of Signaling Lipids. Angewandte Chemie 61, e202205855 (2022). 10.1002/anie.202205855

39 Junge, S. et al. A micro-LED array based platform for spatio-temporal optogenetic control of various cardiac models. Scientific reports 13, 19490 (2023). 10.1038/s41598-023-46149-1

40 Zhou, L. et al. Wireless Self-Powered Optogenetic System for Long-Term Cardiac Neuromodulation to Improve Post-MI Cardiac Remodeling and Malignant Arrhythmia. Adv Sci (Weinh) 10, e2205551 (2023). 10.1002/advs.202205551

41 Yin, R. T. et al. Open thoracic surgical implantation of cardiac pacemakers in rats. Nat Protoc (2022). 10.1038/s41596-022-00770-y

42 Kmecova, J. & Klimas, J. Heart rate correction of the QT duration in rats. European journal of pharmacology 641, 187–192 (2010). 10.1016/j.ejphar.2010.05.038

43 Zacchigna, S. et al. in Cardiovasc Res Vol. 117 43–59 (2021).

44 Derangeon, M. et al. Transforming growth factor beta receptor inhibition prevents ventricular fibrosis in a mouse model of progressive cardiac conduction disease. Cardiovasc Res 113, 464–474 (2017). 10.1093/cvr/cvx026

