## Supplemental figures for "*In vivo* optical induction of I_NaL_ by photoactivation of AaH-II peptide in beating heart using wireless implantable device"

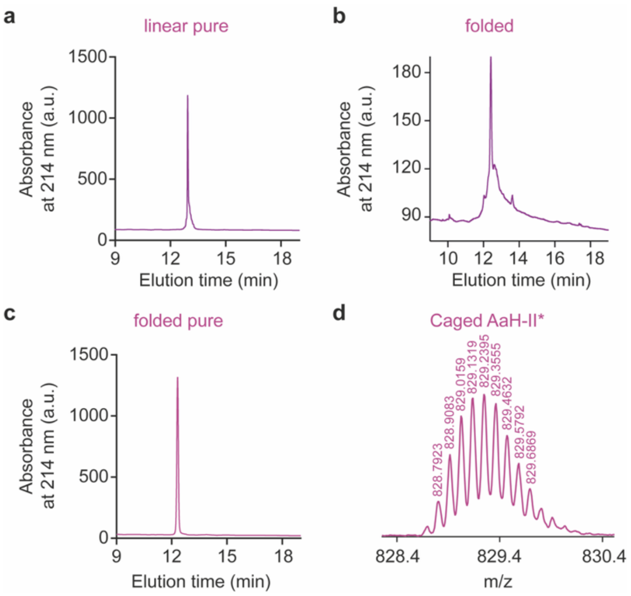


**Supplementary Figure 1. Purification and characterization of caged AaH-II*. (a-c)** RP-HPLC elution profiles of linear purified caged AaH-II* **(a)**, folded/oxidized caged AaH-II* **(b)**, and purified folded/oxidized caged AaH-II* **(c). (d)** LC-ESI QTOF analysis of caged AaH-II* showing an exact mass of 828.7923 $[M+9H]^{9+}$.

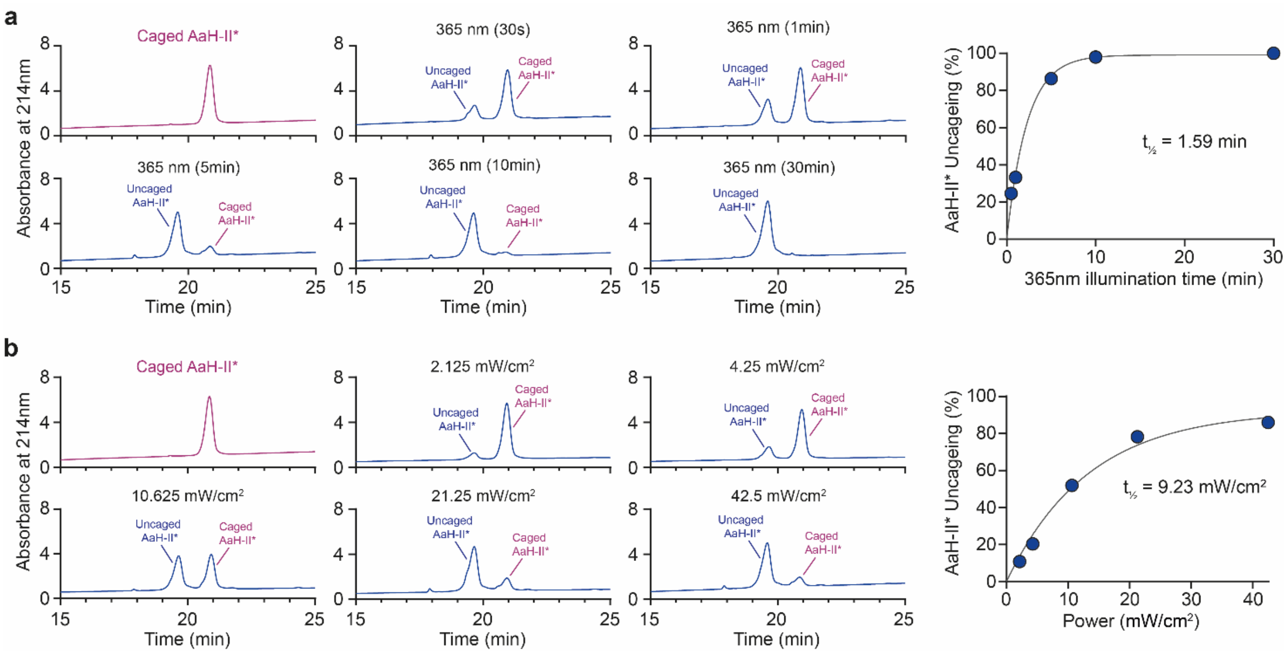


**Supplementary Figure 2. Physicochemical properties of caged AaH-II*. (a)** RP-HPLC profiles of caged AaH-II* for illumination durations at 365 nm and 45 mW/cm$^2$: 0, 30 sec, 1 min, 5 min, 10 min and 30 min **(Left panel)**. Proportion of uncaged AaH-II* quantified from HPLC peak areas as a function of irradiation time at 365 nm **(Right panel)**. **(b)** RP-HPLC profiles of caged AaH-II* with different illumination intensities at 365 nm (5 min duration): 2.12, 4.25, 10.62, 21.25, 42.50 mW/cm$^2$ **(Left panel)**: Proportion of uncaged AaH-II* quantified from HPLC peak areas as a function of irradiation power at 365 nm **(Right panel).**

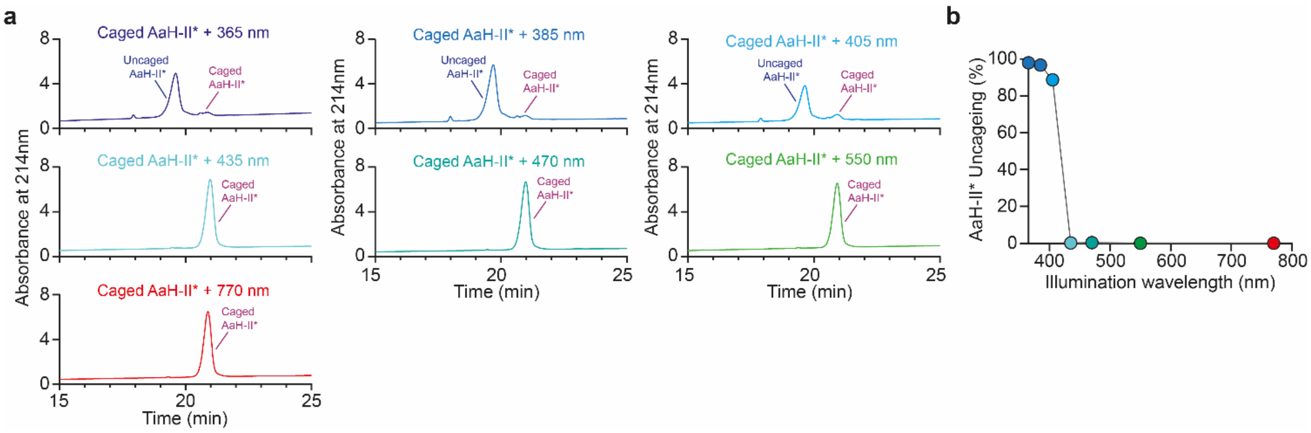


**Supplementary Figure 3. Wavelength-dependent uncaging of AaH-II* (a)** RP-HPLC profiles of caged AaH-II* after 10 min illumination under different wavelengths using constant 45 mW/cm$^2$ irradiation: 365, 385, 405, 435, 470, 550 and 770 nm. **(b)** Proportion of uncaged AaH-II* quantified from HPLC peak areas as a function of the illumination wavelength.

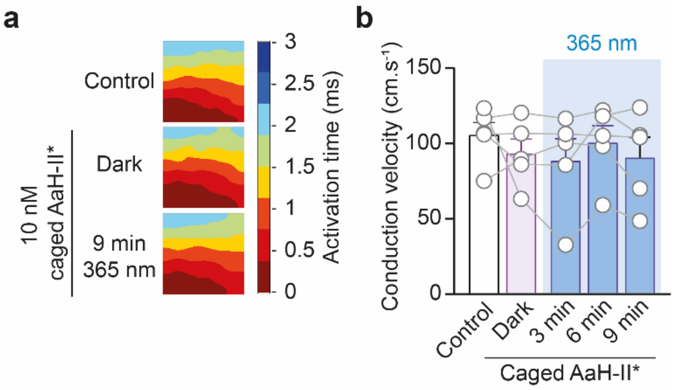


**Supplementary Figure 4. Photoactivation of caged AaH-II* alters ventricular conduction velocity. (a)** Representative MEA activation maps under control conditions, during 10 nM caged AaH-II* perfusion in the dark and after 9 min of 365 nm illumination. **(b)** Corresponding quantification of the conduction velocity. Data are presented as mean ± SEM. Friedman test followed by Dunn's multiple-comparison test *versus* baseline. *, $p < 0.05$.

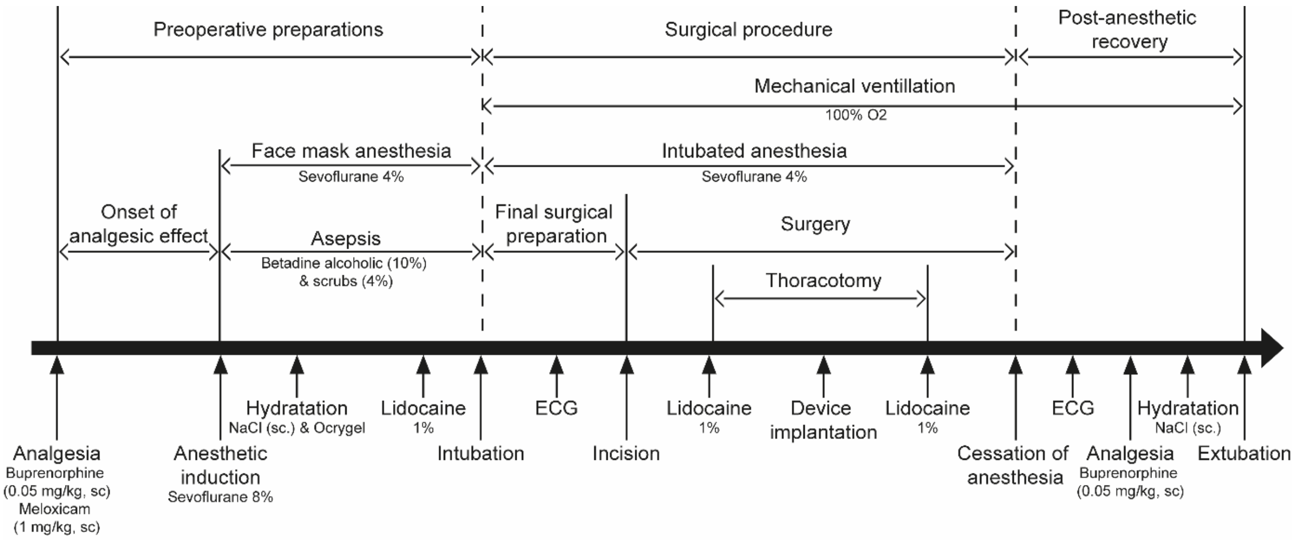


**Supplementary Figure 5. Detailed timeline of the surgical procedure and the peri-operative care.**

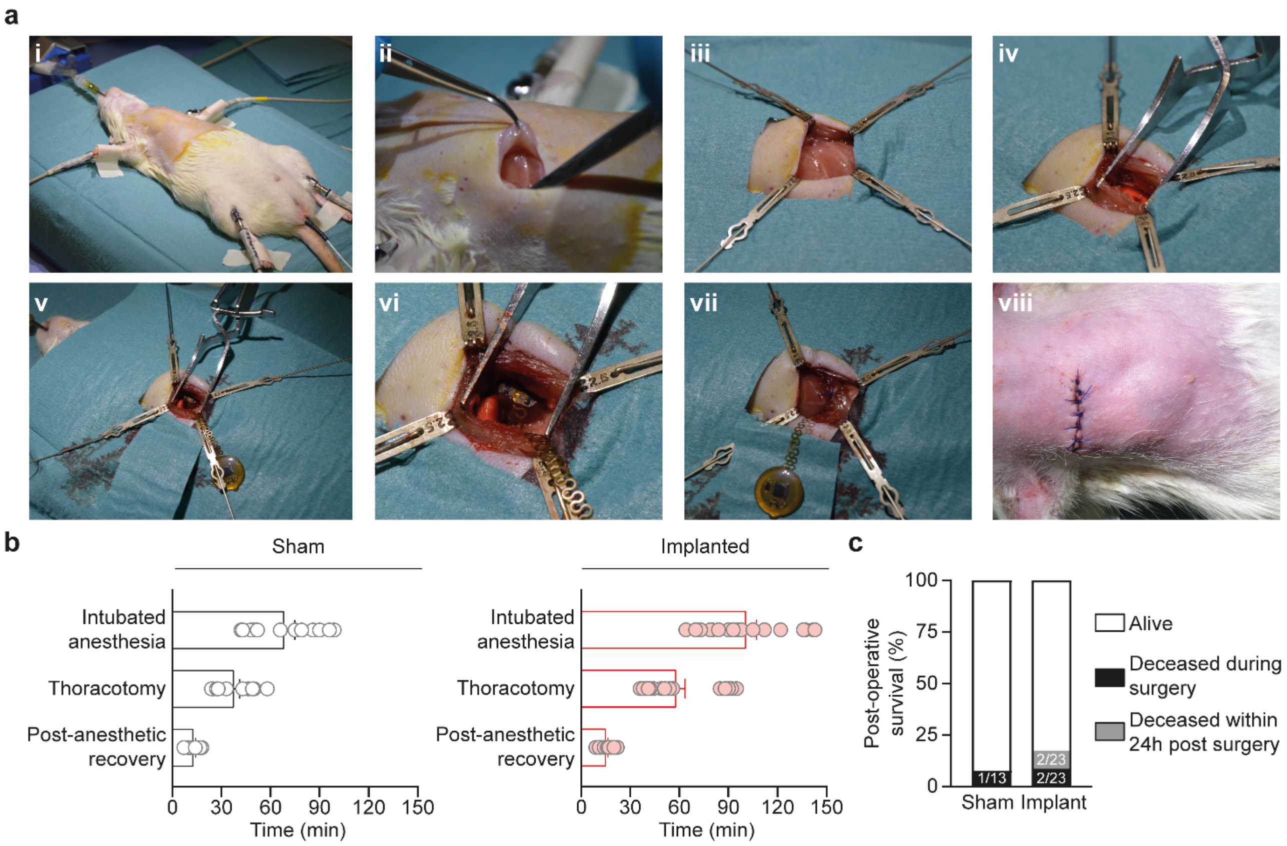


**Supplementary Figure 6. Surgical procedure for device implantation on the right ventricle of rats. (a)** Device implantation in 10-12-week-old adult rats. The surgery is carried out under anesthesia with mechanical ventilation **(i)**. Pectoral muscles are separated **(ii)** and ribs exposed (iii) to access the right lung **(iv)**. The stimulation pad is sutured on the upper part of the RV **(v)** and the thoracic cage is closed, while recreating the pleural vacuum **(vi)**. Muscles are repositioned and the receiver is positioned subcutaneously **(vii)**. (b) Mean duration of the operative phases, including intubated anesthesia, thoracotomy, and post-anesthetic recovery, in sham **(left)** and implanted rats (right). **(c)** 24-hour post-operative survival rate of sham and implanted rats.

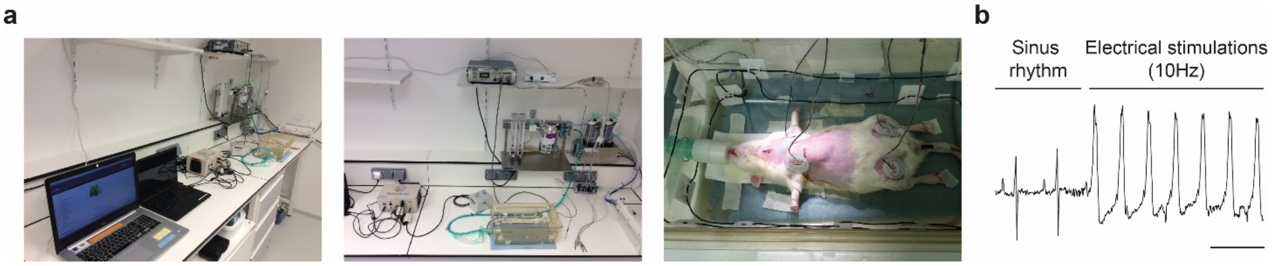


**Supplementary Figure 7. Experimental setup for *in vivo* cardiac photopharmacology. (a)** Experimental setup with the device activation system, ECG recording, gaseous anesthesia and heat monitoring. **(b)** ECG trace during 10-Hz electrical stimulation confirming device function. Scale bar = 200 ms.

**Supplementary Table 1. ECG parameters measured during the *in vivo* activation of caged AaH-II* in anesthetized rats following mini-thoracotomy.**

| Parameters | Open thorax rats with NaCl w/o 365 nm (n=4) | | | | Open thorax rats with caged AaH-II* w/o 365 nm (n=7) | | | |
|---|---|---|---|---|---|---|---|---|
| | Baseline | NaCl + Dark | NaCl + 365nm | NaCl + Dark | Baseline | Caged AaH-II* + Dark | Caged Aa-II* + 365 nm | Caged AaH-II* + Dark |
| RR (ms) | 143.3 ± 5.79 | 142 ± 4.42 | 141.9 ± 2.87 | 143.4 ± 2.5 | 148.2 ± 3.86 | 148.5 ± 3.64 | 144.7 ± 3.39 | 145.7 ± 3.65 |
| PR (ms) | 49.6 ± 2.27 | 49.72 ± 2.24 | 49.88 ± 1.96 | 50.21 ± 1.75 | 52.36 ± 1.33 | 52.33 ± 1.64 | 52.27 ± 1.62 | 51.94 ± 1.51 |
| P duration (ms) | 17.56 ± 1.9 | 17.69 ± 1.86 | 17.75 ± 1.52 | 17.85 ± 1.43 | 17.65 ± 1.17 | 17.39 ± 0.83 | 17.36 ± 1.02 | 17.35 ± 1.03 |
| QRS (ms) | 17.6 ± 0.56 | 17.44 ± 0.62 | 17.4 ± 0.63 | 17.25 ± 0.71 | 18.01 ± 0.64 | 18.13 ± 0.49 | 18.12 ± 0. 53 | 18.16 ± 0.58 |
| QT (ms) | 67.9 ± 1.94 | 69.45 ± 1.39 | 67.92 ± 2.15 | 67.44 ± 1.76 | 64.7 ± 1.01 | 65.19 ± 1.24 | 66.05 ± 1.12* | 65.65 ± 1.26 |
| QTc (ms) | 69.56 ± 2.06 | 71.43 ± 1.31 | 69.87 ± 2.51 | 69.03 ± 2.21 | 65.17 ± 1.096 | 65.61 ± 1.42 | 67.31 ± 1.248** | 66.71 ± 1.537* |
| T Area (mV.ms) | 3.73 ± 0.64 | 3.85 ± 0.7 | 3.62 ± 0.66 | 3.58 ± 0.69 | 3.07 ± 0.52 | 2.94 ± 0.42 | 3.67 ± 0.62 | 3.74 ± 0.61* |
| Normalized T Area | 1 ± 0.17 | 1.033 ± 0.19 | 0.97 ± 0.18 | 0.96 ± 0.18 | 1 ± 0.17 | 0.96 ± 0.14 | 1.2 ± 0.2 | 1.22 ± 0.2* |
| Relative T Area fold change vs. Baseline | 0 | 2.71 ± 2.69 | –3.16 ± 2.49 | –4.57 ± 4.13 | 0 | –0.93 ± 5.82 | 20.4 ± 9.91 | 23.83 ± 10.48 |

Results are presented as mean ± SEM. *, $p<0.05$; **, $p<0.01$; ***, $p<0.001$ indicate statistical differences with baseline group as determined by Friedman test and Dunn's multiple comparisons test.

**Supplementary Table 2. ECG parameters measured in sham and implanted rats before and after the surgery.**

| Parameters | Sham rats (n = 6) | | Implanted rats (n=9) | |
|---|---|---|---|---|
| | Baseline | Post surgery | Baseline | Post surgery |
| RR (ms) | 158.9 ± 6.45 | 159.7 ± 4.51 | 167 ± 6.11 | 174.3 ± 10.34 |
| PR (ms) | 44.88 ± 0.68 | 48.61 ± 1.38* | 49.69 ± 1.53♯ | 50.86 ± 2.52 |
| P duration (ms) | 16.96 ± 0.86 | 19.88 ± 0.51 | 18.3 ± 0.89 | 18.34 ± 0.74 |
| QRS (ms) | 18.53 ± 0.32 | 19.07 ± 0.44 | 20.26 ± 0.68♯ | 19.91 ± 0.51 |
| QT (ms) | 70.42 ± 1.01 | 72.1 ± 1.70 | 68.82 ± 2.111 | 72.77 ± 3.67* |
| QTc (ms) | 68.6 ± 1.55 | 70.03 ± 2.17 | 65.22 ± 1.10 | 67.49 ± 1.75 |
| T Area (mV.ms) | 1.26 ± 0.09 | 2.15 ± 0.55 | 0.87 ± 0.06♯ | 1.14 ± 0.16** |

Results are presented as mean ± SEM. *, p<0.05; **, p<0.01 indicate statistical differences with baseline within a group as determined by Wilcoxon matched-pairs signed-rank test. ♯, p<0.05 indicate statistical difference between the implanted and sham rats as determined by Mann-Whitney test.

**Supplementary Table 3. ECG parameters measured in sham and implanted rats during the 2-week post-operative follow-up.**

| Parameters | Sham rats (n = 6) | | | | Implanted rats (n=5) | | | |
|---|---|---|---|---|---|---|---|---|
| | Baseline | 24 Hrs | 1 Week | 2 Weeks | Baseline | 24 Hrs | 1 Week | 2 Weeks |
| RR (ms) | 158.9 ± 6.45 | 139.9 ± 2.49* | 150.9 ± 4.38 | 155.4 ± 2.92 | 165.4 ± 10.85 | 151.6 ± 3.47* | 156.2 ± 3.86 | 150.5 ± 4.12* |
| PR (ms) | 44.88 ± 0.68 | 44.34 ± 0.89 | 45.56 ± 1.03 | 45.88 ± 1.09 | 52.1 ± 0.82♯♯ | 52.14 ± 1.23♯♯ | 51.8 ± 1.36♯♯ | 49.49 ± 1.12 |
| P duration (ms) | 16.96 ± 0.85 | 16.43 ± 0.73 | 18.9 ± 1.00 | 16.68 ± 0.65 | 17.92 ± 1.30 | 17.69 ± 0.45 | 17.9 ± 0.97 | 18.03 ± 0.45 |
| QRS (ms) | 18.53 ± 0.32 | 19.47 ± 0.24 | 19.42 ± 0.28 | 20.1 ± 0.47 | 19.81 ± 0.53 | 19.34 ± 0.42 | 19.72 ± 0.45 | 20.11 ± 0.62 |
| QT (ms) | 70.42 ± 1.01 | 70.24 ± 1.95 | 69.61 ± 1.41 | 73.35 ± 1.16 | 68.17 ± 3.29 | 71.37 ± 3.14 | 66.76 ± 2.07 | 68.84 ± 2.07 |
| QTc (ms) | 68.6 ± 1.55 | 72.78 ± 2.11 | 69.43 ± 1.00 | 72.06 ± 0.75 | 64.11 ± 1.728 | 69.37 ± 2.392 | 65.03 ± 2.362 | 70.2 ± 1.542 |
| T Area (mV.ms) | 1.26 ± 0.09 | 1.49 ± 0.21 | 1.38 ± 0.20 | 1.56 ± 0.23 | 1.15 ± 0.25 | 2.09 ± 0.31* | 1.36 ± 0.27 | 1.63 ± 0.32 |

Results are presented as mean ± SEM. *, $p<0.05$ indicate statistical differences with baseline within a group as determined by Friedman test and Dunn's multiple comparisons test. ♯♯, $p<0.01$ indicate statistical difference between the implanted and sham rats as determined by Mann-Whitney test.

**Supplementary Table 4. Echocardiographic parameters measured in sham and implanted rats during the 2-week post-operative follow-up.**

| Parameters | Sham rats | | | Implanted rats | | |
|---|---|---|---|---|---|---|
| | Baseline | 2 weeks | | Baseline | 2 weeks | |
| E/A (mm) | 1.28 ± 0.04 | 1.19 ± 0.04 | (n = 5) | 1.42 ± 0.08 | 1.25 ± 0.04 | (n = 6) |
| LVEF (%) | 81.82 ± 2.17 | 82.28 ± 1.49 | (n = 10) | 87 ± 2.39 | 86.83 ± 1.75 | (n = 6) |
| TAPSE (mm) | 2.98 ± 0.11 | 2.52 ± 0.12* | (n = 10) | 3.18 ± 0.37 | 3.36 ± 0.18♯♯♯ | (n = 6) |
| TEP (mm) | 67.14 ± 2.68 | 62.58 ± 3.89 | (n = 10) | 79.67 ± 3.35♯♯ | 81.5 ± 3.71♯♯ | (n = 6) |
| TAP (mm) | 27.33 ± 4.06 | 22.33 ± 4.01 | (n = 8) | 33.5 ± 2.84 | 33.33 ± 3.93 | (n = 6) |

Results are presented as mean ± SEM. *, p<0.05 and **, p<0.01 indicate statistical differences from baseline within a group as determined by Wilcoxon matched-pairs signed-rank test. ♯♯, p<0.01 and ♯♯♯, p<0.001 indicate statistical differences between the implanted and sham rats as determined by Mann-Whitney test. E/A: early/late diastolic transmitral flow velocity; LVEF: left ventricular ejection fraction; TAPSE: tricuspid annular plane systolic excursion; TEP: total ejection period; TAP: time to aortic peak.

**Supplementary Table 5. ECG parameters measured implanted rats with 405 nm device mediated illumination.**

| | Implanted rats with 405 nm (n=8) | |
|---|---|---|
| Parameters | Baseline | Light ON |
| RR (ms) | 148.8 ± 3.88 | 148.8 ± 3.85 |
| PP (ms) | 148.8 ± 3.90 | 148.7 ± 3.87 |
| PR (ms) | 50.55 ± 1.98 | 50.41 ± 1.95 |
| P duration (ms) | 16.55 ± 0.94 | 16.56 ± 0.88 |
| QRS (ms) | 20.78 ± 0.58 | 20.76 ± 0.63 |
| QT (ms) | 72.96 ± 3.13 | 72.82 ± 3.29 |
| QTc (ms) | 73.27 ± 2.96 | 73.15 ± 3.12 |
| T Area (mV.ms) | 2.2 ± 0.37 | 2.2 ± 0.38 |
| Normalized T Area | 1 ± 0.17 | 1 ± 0.17 |
| Relative T Area fold change vs. Baseline | 0 | –0.093 ± 0.99 |

Results are presented as mean ± SEM. No statistical differences have been reported using Wilcoxon matched-pairs signed-rank test.

**Supplementary Table 6. ECG parameters measured implanted rats injected with 67 µg/kg caged AaH-II* with or without the 405-nm device-mediated illumination.**

| | Implanted rats with 67 µg/kg caged AaH-II* (n=5) | | | | Implanted rats with 67 µg/kg caged AaH-II* and 405 nm (n=8) | | | |
|---|---|---|---|---|---|---|---|---|
| Parameters | Baseline | Caged AaH-II* 3 min | Caged AaH-II* 13 min | Caged AaH-II* 23 min | Baseline | Caged AaH-II* 3 min | Caged AaH-II* 13 min + Light ON | Caged AaH-II* 23 min + Light OFF |
| RR (ms) | 151.4 ± 4.89 | 154.3 ± 7.01 | 149.3 ± 3.41 | 147.5 ± 3.41 | 148.8 ± 3.88 | 163.2 ± 5.64 | 156.3 ± 5.74 | 160.6 ± 11.70 |
| PP (ms) | 151.4 ± 4.89 | 154.3 ± 7.01 | 149.3 ± 3.41 | 147.5 ± 3.41 | 148.8 ± 3.90 | 163.1 ± 5.71 | 156.3 ± 5.73 | 160.6 ± 11.93 |
| PR (ms) | 48.39 ± 2.99 | 49.16 ± 2.87 | 49.44 ± 2.68 | 49.89 ± 2.85 | 50.55 ± 1.98 | 51.13 ± 2.06 | 52.01 ± 2.27 | 54.63 ± 3.88 |
| P duration (ms) | 14.91 ± 0.93 | 14.47 ± 1.19 | 14.89 ± 0.96 | 14.96 ± 0.97 | 16.55 ± 0.94 | 16.21 ± 0.69 | 16.19 ± 0.54 | 16.67 ± 0.69 |
| QRS (ms) | 19.46 ± 0.76 | 19.47 ± 0.83 | 19.25 ± 0.58 | 19.29 ± 0.70 | 20.78 ± 0.58 | 21.13 ± 0.67 | 21.27 ± 0.83 | 21.3 ± 0.87 |
| QT (ms) | 74.3 ± 6.43 | 76.83 ± 6.36 | 74.32 ± 6.90 | 74.25 ± 7.56 | 72.96 ± 3.13 | 76.18 ± 4.61 | 78.2 ± 5.70 | 76.17 ± 4.56 |
| QTc (ms) | 73.83 ± 5.99 | 75.54 ± 5.33 | 74.28 ± 6.32 | 74.79 ± 7.50 | 73.27 ± 2.96 | 72.94 ± 3.73 | 76.28 ± 4.36 | 73.82 ± 3.41 |
| T Area (mV.ms) | 3.33 ± 0.34 | 3.74 ± 0.44 | 3.16 ± 0.27 | 3.17 ± 0.38 | 2.195 ± 0.37 | 2 ± 0.44 | 3.11 ± 0.83** | 2.85 ± 0.41*** |
| Normalized T Area | 1 ± 0.10 | 1.12 ± 0.13 | 0.95 ± 0.082 | 0.95 ± 0.12 | 1 ± 0.17 | 0.91 ± 0.20 | 1.417 ± 0.38** | 1.299 ± 0.19*** |
| Relative T Area fold change vs. Baseline | 0 | 11.93 ± 3.68 | –4.52 ± 3.13 | –5.36 ± 7.10 | 0 | –12.21 ± 4.38 | 35.04 ± 21.93 | 35.11 ± 13.33 |

Results are presented as mean ± SEM. **, p<0.01 and ***, p<0.001 indicate statistical differences with the caged AaH-II* 3 min condition as determined by Friedman test and Dunn's multiple comparisons test
